# Axonal transport of Hrs is activity-dependent and rate limiting for synaptic vesicle protein degradation

**DOI:** 10.1101/2020.04.16.044818

**Authors:** Veronica Birdsall, Mei Zhu, Konner Kirwan, Yuuta Imoto, Shigeki Watanabe, Clarissa L. Waites

## Abstract

Turnover of synaptic vesicle (SV) proteins is vital for the maintenance of healthy and functional synapses. SV protein turnover is driven by neuronal activity in an ESCRT (endosomal sorting complex required for transport)-dependent manner. Here, we characterize a critical step in this process: axonal transport of ESCRT-0 component Hrs, necessary for sorting proteins into the ESCRT pathway and recruiting downstream ESCRT machinery to catalyze multivesicular body (MVB) formation. We find that neuronal activity stimulates the formation of presynaptic endosomes and MVBs, as well as the motility of Hrs+ vesicles in axons and their delivery to SV pools. Hrs+ vesicles co-transport ESCRT-0 component STAM1 and comprise a subset of Rab5+ vesicles, likely representing pro-degradative early endosomes. Furthermore, we identify kinesin motor protein KIF13A as essential for the activity-dependent transport of Hrs to SV pools and the degradation of SV membrane proteins. Together, these data demonstrate a novel activity- and KIF13A-dependent mechanism for mobilizing axonal transport of ESCRT machinery to facilitate the degradation of SV membrane proteins.

## Introduction

Synaptic vesicles (SVs) are the fundamental units of neurotransmitter release, and as such their efficient fusion and recycling are essential for neuronal communication. These processes depend upon maintaining the proper complement of functional SV membrane proteins. Indeed, the inability to clear damaged or misfolded SV proteins can precipitate synaptic dysfunction and neurodegeneration (Bezprozvanny and Hiesinger 2013; Esposito, Ana Clara, and Verstreken 2012; Hall et al. 2017). However, the elaborate morphology of neurons creates unique spatial challenges for SV protein clearance and degradation. First, lysosomes - the degradative organelles for SV membrane proteins - are primarily localized to neuronal cell bodies, requiring retrograde transport of SV proteins destined for degradation. Second, the machinery responsible for packaging and delivering SV proteins to lysosomes must be transported from the cell body to presynaptic boutons, which can be many meters away. In addition to these spatial challenges, neurons face temporal challenges in transporting degradative machinery in response to stimuli such as synaptic activity. For instance, degradative organelles (proteasomes, lysosomes, and autophagosomes) undergo activity-dependent recruitment into the postsynaptic compartment as part of the mechanism for synaptic plasticity (Bingol and Schuman 2006; Goo et al. 2017; Shehata et al. 2012), and must be rapidly mobilized to these sites. Similarly, the turnover of SV and other presynaptic proteins is stimulated by neuronal activity (Sheehan et al. 2016; Truckenbrodt et al. 2018), requiring the local presence of degradative machinery to facilitate clearance of old and damaged proteins. However, very little is known about whether or how neurons regulate the transport and delivery of this machinery to presynaptic terminals.

Previous work from our group and others has demonstrated that the degradation of SV membrane proteins requires the endosomal sorting complex required for transport (ESCRT) pathway (Sheehan et al. 2016; Uytterhoeven et al. 2011). Comprising a series of protein complexes (ESCRT-0, -I, -II, -III, Vps4), the ESCRT pathway recruits ubiquitinated membrane proteins and forms multivesicular bodies (MVBs) for delivery of this cargo to lysosomes (Hurley 2015). In addition to its degradative function, the ESCRT pathway also functions in other cellular processes, including cytokinesis and plasma membrane repair (Stoten and Carlton 2018; McCullough, Frost, and Sundquist 2018; Christ et al. 2017; Jimenez et al. 2014). Mutation or dysfunction of ESCRT-associated proteins leads to neurodegeneration in animal models (Tamai et al. 2008; J. A. Lee et al. 2007; D. Lee et al. 2019) and is linked to frontotemporal dementia (FTD)(Skibinski et al. 2005; J. A. Lee et al. 2007), amyotrophic lateral sclerosis (ALS)(Parkinson et al. 2006) and infantile-lethal epilepsy (Hall et al. 2017) in humans. Moreover, the ESCRT pathway is essential for endosomal protein trafficking and implicated in the etiology of Alzheimer’s and Parkinson’s diseases (Vaz-Silva et al. 2018; Borland and Vilhardt 2017; Vagnozzi and Praticò 2019; Feng et al. 2020; Spencer et al. 2014, 2016). Yet despite the essential role of the ESCRT pathway in protein degradation and other cellular processes, as well as its links to neurodegenerative disease, little is known about its substrates, dynamic behavior, or mechanisms of localization and regulation in neurons.

Our previous study demonstrated that a subset of SV membrane proteins undergo activity-dependent degradation in a process initiated by the activation of small GTPase Rab35 and the recruitment of its effector, ESCRT-0 component Hrs (Sheehan et al. 2016). As the first component of the ESCRT pathway, Hrs functions to sequester ubiquitinated proteins destined for degradation, and to recruit downstream ESCRT machinery to promote MVB biogenesis. Interestingly, neuronal activity significantly increases the localization of Hrs to axons and SV pools, suggesting that it may be transported to SV pools in an activity-dependent manner to catalyze SV protein degradation. However, it is not clear how Hrs is transported into and out of synapses.

In the current study, we use live imaging to characterize Hrs localization and transport in axons. We find that Hrs is transported on vesicles that exhibit activity-induced anterograde and bidirectional motility. These vesicles carry ESCRT-0 protein STAM1 and comprise a subset of Rab5+ early endosomes in axons. In addition, we identify the kinesin motor protein KIF13A as essential for the activity-dependent transport of Hrs; not only does neuronal activity stimulate the binding of Hrs and KIF13A, but KIF13A knockdown inhibits the activity-induced trafficking of Hrs along axons and its delivery to SV pools, as well as SV protein degradation. Together, these findings illuminate critical mechanisms of ESCRT-0 protein transport and delivery to presynaptic boutons in order to facilitate SV protein degradation.

## Results

### Neuronal activity increases the density of presynaptic endosomes

In our previous work, we found that neuronal firing increased the localization of ESCRT-0 protein Hrs to axons and SV pools (Sheehan et al. 2016). Given the role of Hrs in recruiting downstream ESCRT machinery to catalyze MVB formation (Raiborg, Rusten, and Stenmark 2003), we hypothesized that Hrs recruitment would serve to increase the number of MVBs and other endosomal structures at presynaptic boutons. We therefore examined electron micrographs of boutons from 16-21 day *in vitro* (DIV) hippocampal neurons treated for 48 hours with tetrodotoxin (TTX) to reduce neuronal firing or gabazine to increase firing (Fig. 1A, B). Indeed, we observed a significantly higher density of presynaptic MVBs and endosomes (defined as single-membrane structures >80 nm in diameter; Fig. 1B, C) following gabazine versus TTX treatment (Fig. 1A, E), suggesting that neuronal activity promotes the appearance of these structures at boutons. Since other studies have reported that neuronal firing stimulates the biogenesis of autophagosomes at presynaptic terminals (Wang et al. 2015), we also counted the number of multilamellar structures present under each condition (Fig. 1D, F). Interestingly, not only was the average density of these structures lower than that of endosomes/MVBs regardless of activity level (∼0.1 multilamellar structures/μm^2^ vs. ∼1 endosomes/μm^2^ at baseline), but their number did not increase with gabazine treatment (Fig. 1F), indicating that prolonged neuronal activity specifically increases the number of endosomes and MVBs at presynaptic boutons (Sheehan et al. 2016). To examine whether Hrs associates with such structures, we performed correlative-light electron microscopy (CLEM) in cultured hippocampal neurons co-expressing HaloTag-Hrs and vesicular glutamate transporter (vGlut1-GFP) to label SV pools. Indeed, we found Hrs labeling associated with endosomal structures including an MVB (Fig. 1G, arrows), confirming its presence at putative sites of endolysosomal sorting.

**Figure 1.**
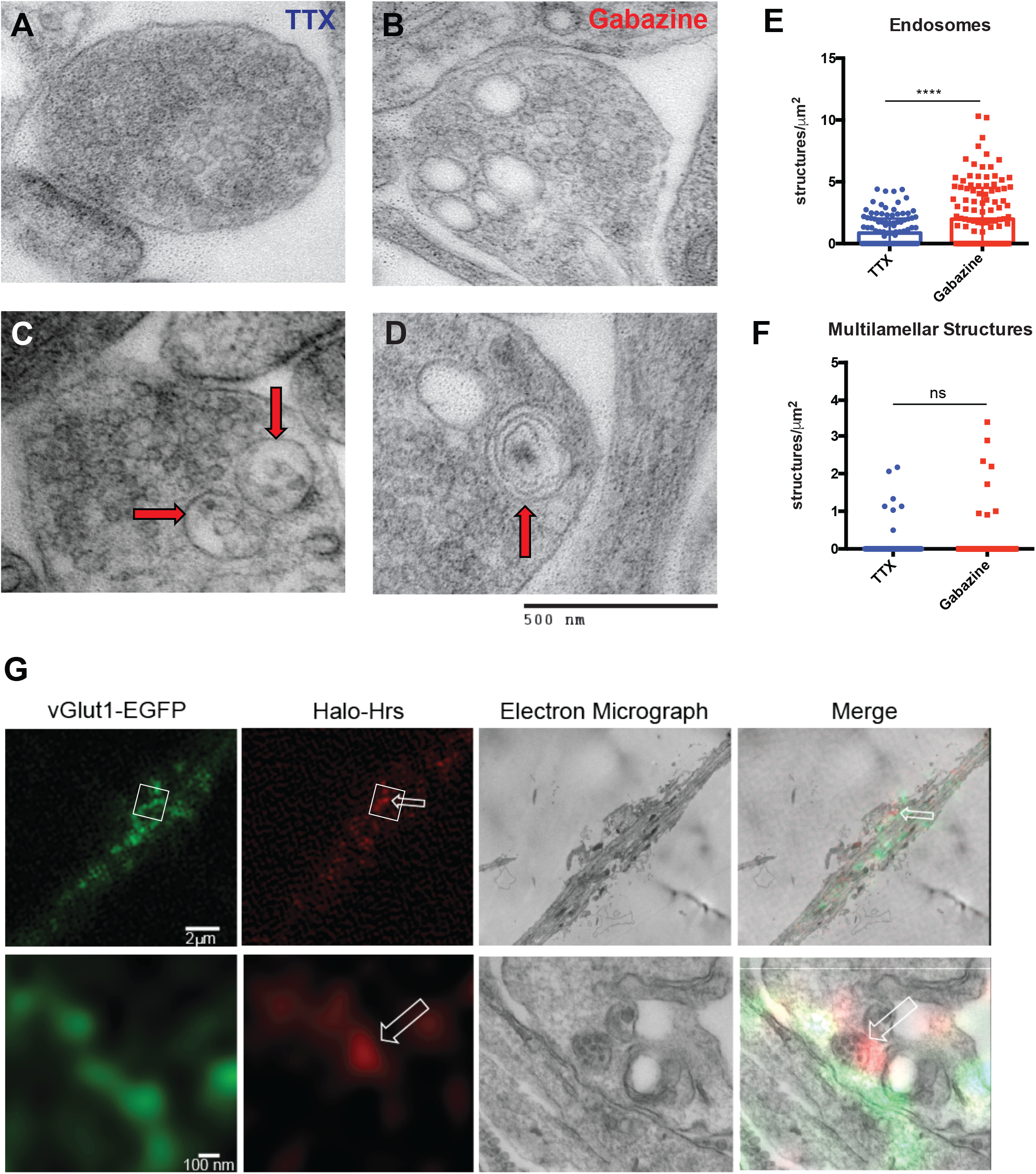
Neuronal activity increases the number of presynaptic endosomes and Hrs associates with these structures. **(A)** Representative electron micrograph of a presynaptic bouton from hippocampal neurons treated with TTX for 48hrs. **(B)** Representative electron micrograph of a presynaptic bouton from hippocampal neurons treated with gabazine for 48h. **(C- D)** Electron micrographs of structures scored as endosomal (single-membrane, >80nm diameter, red arrows; **C**) and multilamellar (>1 membrane, red arrows; **D**). Size bar, 500 nm. **(E-F)** Quantification of endosomal **(E)** and multilamellar **(F)** structures per μm^2^ of presynaptic area at TTX- or gabazine-treated boutons. The density of endosomal structures, but not multilamellar structures, is significantly increased following gabazine treatment. Scatter plot shows median and interquartile range (****P<0.0001, ns P=0.8191, Mann-Whitney test; n=102 (TTX-treated) and 106 (gabazine-treated) boutons/condition). **(G)** Representative CLEM images of axons from hippocampal neurons expressing vGlut1-EGFP (green) and Halo-Hrs (Jenelia-Halo-549 nm; red), treated for 2hrs with Bic/4AP (upper row: lower magnification images of putative axon; lower row: higher magnification images of Halo-Hrs-positive MVB (white arrows) in white boxed area. Size bars were labeled as indicated in the images.

### Neuronal activity stimulates the anterograde and bidirectional transport of Hrs

These ultrastructural findings, together with our previous work (Sheehan et al. 2016), suggest that the localization and transport of Hrs are regulated by neuronal activity to catalyze SV protein degradation. We thus investigated whether fluorescently-tagged Hrs exhibited activity-dependent changes in motility or directional transport in axons. To verify that overexpressed Hrs reflected the behavior of the endogenous protein, we immunostained 14-15 day *in vitro* (DIV) hippocampal neurons treated with DMSO (control) or 40 μm bicuculline and 50 μm 4-aminopyridine (Bic/4AP) for 20 hours. Endogenous Hrs exhibited a punctate distribution along axons (Fig. S1A), similar to the fluorescently-tagged protein (Sheehan et al. 2016). Moreover, Bic/4AP treatment led to an increase in both the intensity and number of Hrs puncta per unit length axon compared to the control condition (Fig. S1B, C), consistent with our previous findings with RFP-Hrs.

We next performed imaging experiments in tripartite microfluidic chambers that isolate cell bodies from axons while enabling synapse formation between neurons in adjacent compartments (Taylor et al. 2005; Park et al. 2006; Batista, Martínez, and Hengst 2017) (Fig. S2A). Lentiviral transduction of cell bodies in these chambers led to moderate protein expression levels and consistent transduction efficiency across constructs used in this study (∼50%; Fig. S2B). Imaging of RFP-Hrs in 13-15 DIV neurons was performed either immediately following application of DMSO (control) or Bic/4AP (acute Bic/4AP), or 2 hours after treatment with these drugs (2h Bic/4AP)(Fig. 2A-C; Videos 1&2), which did not cause any detectable changes in neuronal health over this timeframe (Fig. S2C). Under control conditions, we observed that >60% of Hrs puncta were stationary over the course of the ∼4-minute imaging session (Fig. 2A, D, F). Remarkably, acute treatment with Bic/4AP led to a slight but significant increase in puncta motility (from ∼38% to ∼50% motile), with the effect growing stronger after 2h of treatment (∼60% motile puncta; Fig. 2A-D, F). No change in speed was observed across the three conditions (Fig. 2E), indicating that activity increases the number of Hrs puncta in the motile pool rather than changing the mechanism of transport. Interestingly, inhibition of neuronal firing by TTX did not decrease Hrs motility (Fig. S2D), suggesting that basal activity levels in these cultures are low. Similar effects of Bic/4AP on axonal EGFP-Hrs motility were observed in dissociated hippocampal cultures (Fig. S3A-E), demonstrating that these activity-induced motility changes are not artifacts of imaging in microfluidic chambers.

**Figure 2.**
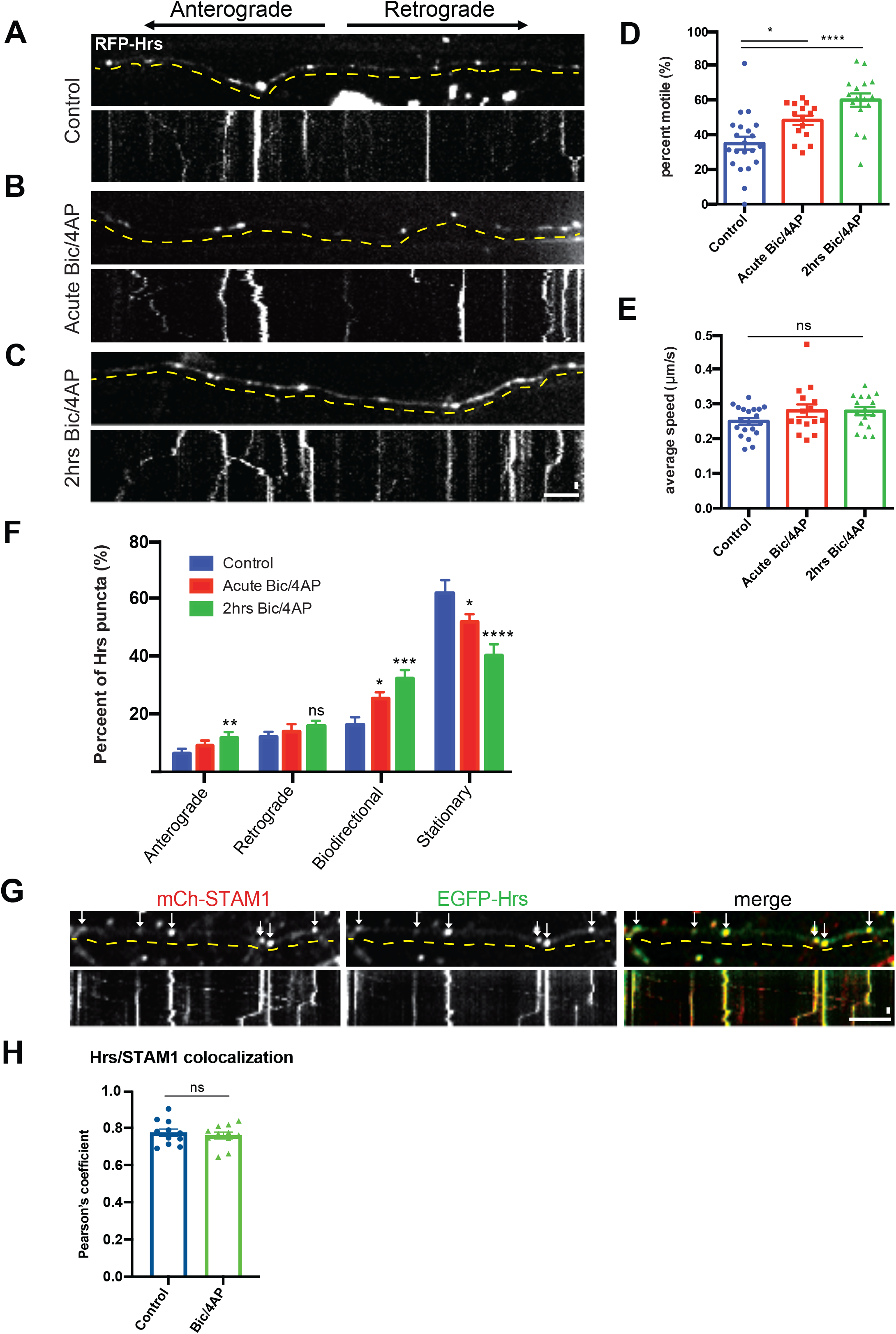
Neuronal activity stimulates the transport of Hrs+ vesicles in axons. **(A-C)** Representative images and corresponding kymographs of RFP-Hrs in microfluidically isolated axons from 13-15 DIV hippocampal neurons under control conditions (**A**; also see Video 1), acute treatment with Bicuculline/4AP (**B**), and after 2hrs treatment with Bicuculline/4AP (**C**; Video 2). **(D)** Percentage of motile Hrs puncta in axons, showing a significant increase with acute and 2hrs Bic/4AP treatment (****P<0.0001, *P=0.0198, one-way ANOVA with Dunnett’s multiple comparisons test; ≥3 separate experiments, n=20 (control), 15 (acute Bic/4AP), 16 (2hrs Bic/4AP) videos/condition). **(E)** Average speed of motile Hrs puncta (μm/sec), showing no change across conditions (ns P=0.1594; same ‘n’ values as D). **(F)** Breakdown of directional movement of Hrs puncta, expressed as percentage of total Hrs puncta. Anterograde and bidirectional movement are significantly increased following 2hrs Bic/4AP treatment (****P<0.0001, ***P=0.0001, **P=0.0064, *P=0.0355 (bidirectional), *P=0.0198 (stationary), ns P=0.3102, one-way ANOVA with Dunnett’s multiple comparisons test; same ‘n’ values as D). **(G)** Representative images and corresponding kymographs for EGFP-Hrs and mCh-STAM1 in axons from dissociated hippocampal neurons under control conditions (also see Video 3). Arrows show Hrs puncta that colocalize with STAM1. **(H)** Pearson’s coefficient of colocalization for EGFP-Hrs and mCh-STAM1, showing no change with 2hrs Bic/4AP treatment (ns P=0.5936, unpaired t-test, n=11 axons from 5 videos/condition). For all images, horizontal size bar =10 µm, vertical size bar =25 s, 1 frame taken every 5 s (0.2 fps). Dashed yellow lines highlight axons. All scatter plots show mean ± SEM.

Like other early endosome-associated proteins (*e.g.* Rab5)(Goto-Silva et al. 2019), Hrs exhibits a mixture of anterograde, retrograde, and non-processive bidirectional motility (Fig. 2F, S3F). However, we found that only anterograde and bidirectional motility increased following treatment with Bic/4AP (Fig. 2F), and that the total net displacement of Hrs puncta was shifted in the anterograde direction after 2h of Bic/4AP (Fig. S3G-H), suggesting that neuronal activity preferentially mobilizes Hrs towards distal axonal sites. Hrs is known to recruit STAM1, the other ESCRT-0 component, in order to form heterotetrameric ESCRT-0 complexes (Mayers et al. 2011; Takahashi et al. 2015). Therefore, we next investigated whether STAM1 was also present on Hrs+ puncta. Co-expression of mCh-STAM1 with EGFP-Hrs in dissociated hippocampal neurons revealed that these proteins were highly colocalized in axons including on motile vesicles (∼80% colocalization; Fig. 2G-H, Video 3) as well as in the somatodendritic compartment (data not shown). To ensure that Hrs overexpression did not influence STAM1’s localization and dynamic behavior, we also evaluated mCh-STAM1 motility in singly-transduced axons in microfluidic chambers. STAM1 dynamics were very similar to those of Hrs, with increased motility after 2h of Bic/4AP treatment, no change in average speed, and an activity-induced increase in the fraction of anterograde and bidirectional puncta (Fig. S4). Together, these data show that the transport of both ESCRT-0 proteins is sensitive to neuronal firing.

### Hrs+ vesicles comprise a subset of Rab5+ vesicles in axons

ESCRT-0 proteins localize to early endosomes (Sun et al. 2003; Raiborg, Rusten, and Stenmark 2003; Flores-Rodriguez et al. 2015; Raiborg et al. 2001), and studies in non-neuronal cells show that Hrs recruitment to endosomal membranes depends upon interactions between its FYVE domain and the endosome-enriched lipid phosphoinositol-3-phosphate (PI3P)(Komada and Soriano 1999; Stahelin et al. 2002). To determine whether this FYVE/PI3P interaction was similarly necessary for Hrs recruitment to axonal vesicles, we treated hippocampal neurons for 24h with an inhibitor of PI3P synthesis, SAR405 (Schu et al. 1993), and examined the number of EGFP-Hrs puncta per unit length axon as well as their fluorescence intensity. SAR405 treatment dramatically decreased both the density (∼39%) and intensity (∼36%) of EGFP-Hrs puncta (Fig. S5A-C), indicating that PI3P is essential for Hrs association with axonal vesicles. This effect was replicated by an inactivating mutation of the Hrs FYVE domain (R183A; Raiborg et al. 2001)(Fig. S5A-C), further confirming that the FYVE/PI3P interaction is critical for Hrs targeting to vesicles. To evaluate whether the axonal dynamics of the FYVE domain recapitulated those of full-length Hrs, we assessed the dynamic behavior of EGFP-2xFYVE in neurons cultured in microfluidic chambers under control, acute Bic/4AP, and 2h Bic/4AP conditions. Interestingly, while the 2xFYVE domain exhibited highly punctate expression in axons, its dynamics were different from those of full-length Hrs and it did not demonstrate activity-dependent changes in motility (Fig. S5D-G), indicating that other domains of Hrs are required for its activity-dependent transport.

We next investigated whether activity-dependent motility is a general feature of early endosomes in axons. Here, we examined the motility of the small GTPase and early endosome marker Rab5, critical for formation and function of the endolysosomal network (Gorvel et al. 1991; Deinhardt et al. 2006; Poteryaev et al. 2010), in axons from microfluidic chambers. We found that Rab5 puncta exhibited higher motility under control conditions than Hrs puncta (∼60% motile puncta at baseline; Fig. 3A, D, Video 4), with no increase in motility upon acute Bic/4AP application (Fig. 3B, D) but a ∼20% increase following 2h of treatment (Fig. 3C, D, Video 5). Similar to Hrs and STAM1, the average speed of Rab5 puncta did not change in response to neuronal firing (Fig. 3E), but the proportion of anterogradely moving puncta increased significantly (Fig. 3F). These data show that while axonal Rab5+ vesicles do exhibit activity-dependent changes in motility, this effect is more pronounced for ESCRT-0+ vesicles.

**Figure 3.**
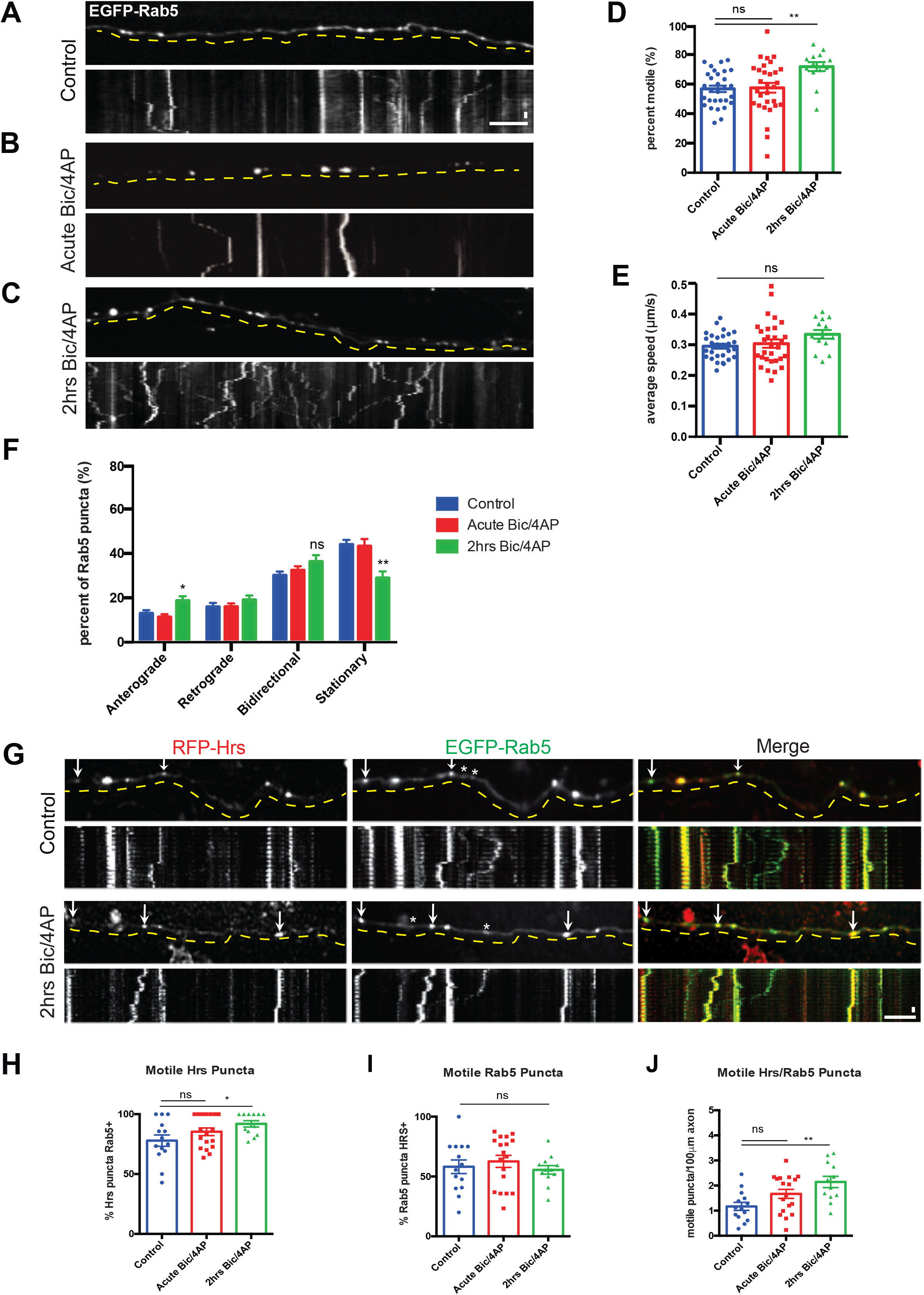
Hrs is present on a subset of Rab5+ vesicles. **(A-C)** Representative images and corresponding kymographs of EGFP-Rab5 in microfluidically isolated axons from 13-15 DIV hippocampal neurons under control (**A**, see Video 4), acute Bic/4AP (**B**), and 2hrs Bic/4AP (**C**, see Video 5) conditions. **(D)** Percentage of motile Rab5 puncta in axons, showing a significant increase only after 2hrs Bic/4AP treatment (**P=0.0013, ns P>0.9999, Kruskal-Wallis test with Dunn’s multiple comparisons; ≥3 separate experiments, n=31 (control), 30 (acute Bic/4AP), 14 (2hrs Bic/4AP) videos/condition). **(E)** Average speed of motile Rab5 puncta (μm/sec) is unchanged across conditions (ns P=0.1295, Same ‘n’ values as D). **(F)** Breakdown of directional Rab5 puncta movement, showing that only anterograde movement is significantly increased following 2hrs Bic/4AP treatment (**P=0.004, *P=0.0404, ns P=0.1082, one-way ANOVA with Dunnett’s multiple comparisons test; same ‘n’ values as D). **(G)** Representative images and corresponding kymographs for RFP-Hrs and EGFP-Rab5 in axons from dissociated hippocampal neurons under control and 2hrs Bic/4AP conditions (see Videos 6&7). Arrows indicate motile puncta that are positive for Hrs and Rab5; stars indicate motile puncta that are only positive for Rab5. **(H)** Percentage of motile Hrs puncta that are Rab5+, showing a significant increase in motility after 2hrs Bic/4AP treatment (*P=0.028, ns P=0.2471, Kruskal-Wallis test with Dunn’s multiple comparisons; ≥3 separate experiments, n=14 (control), 18 (acute Bic/4AP), 12 (2hrs Bic/4AP) videos/condition). **(I)** Percentage of motile Rab5 puncta that are Hrs+, showing no significant difference in motility across conditions (ns P=0.5938, same ‘n’ values as H). **(J)** Percentage of motile puncta that are Hrs+ and Rab5+ per 100 µm of axon, showing a significance increase in motility after 2hrs Bic/4AP treatment (**P=0.0025, ns P=0.1023, one-way ANOVA with Dunnett’s multiple comparisons test; same ‘n’ values as H). For all images, horizontal size bar =10 µm, vertical size bar =25 s, 1 frame taken every 5 s (0.2 fps). Dashed yellow lines highlight axons. All scatter plots show mean ± SEM.

Since the axonal dynamics of Rab5 do not precisely align with those of ESCRT-0 proteins, we next investigated whether Hrs is present on the same vesicles as Rab5. Here, we performed time-lapse imaging of axons from dissociated neurons co-transfected with RFP-Hrs and EGFP- Rab5 (Fig. 3G, Videos 6&7). The density of Rab5 puncta in co-transfected axons was on average higher than that of Hrs puncta, and while >80% of motile Hrs puncta were also Rab5+ (Fig. 3H), the converse was not true. Indeed, only ∼60% of motile Rab5 puncta were Hrs+ (Fig. 3I). These findings demonstrate that Hrs is present on a subset of Rab5+ structures, consistent with previous studies showing that Rab5 associates with a heterogenous population of endosomes as well as SVs (Simonsen et al. 1998; Christoforidis et al. 1999; Flores-Rodriguez et al. 2015; Fischer von Mollard et al. 1994; Shimizu, Kawamura, and Ozaki 2003; Zhang et al. 2013). However, the number of motile Hrs+/Rab5+ puncta increased significantly after 2h Bic/4AP (Fig. 3J), indicating that the Hrs+ subset of Rab5 vesicles is responsive to neuronal firing, even though the total population of Rab5 vesicles shows a more modest increase in activity-induced motility (Fig. 3D).

Given that Rab35 mediates the recruitment of Hrs to SV pools and can colocalize with Hrs on axonal vesicles (Sheehan et al. 2016), we next investigated whether Rab35 transport dynamics were similar to those of Hrs. As with STAM1 and Rab5, we performed timelapse imaging of EGFP-Rab35 in singly-transduced axons in microfluidic chambers. Like Rab5, Rab35 puncta exhibited higher motility than Hrs at baseline; unlike Rab5 and ESCRT-0 proteins, these puncta did not exhibit any changes in motility or directional movement following acute or 2h Bic/4AP treatment (Fig. S6A-E). To determine whether the colocalization of Hrs with Rab35 was affected by neuronal activity, dissociated cultures co-expressing RFP-Hrs and EGFP-Rab35 were treated for 2h with vehicle or Bic/4AP prior to fixation and analysis. We found that Hrs colocalization with Rab35 increased significantly following Bic/4AP treatment (Fig. S6F), suggesting an activity-dependent recruitment of Hrs to Rab35+ puncta. By measuring the colocalization of mCh-Rab35 with EGFP-Synapsin, an SV-associated protein, we found that Rab35 was highly colocalized with SV pools (∼80%, Fig. S6G, H). Moreover, this colocalization was unaffected by 2h Bic/4AP treatment (Fig. S6H), indicating that the activity-induced increase in Hrs/Rab35 colocalization represents Hrs recruitment to SV pools.

### KIF13A interacts with Hrs in an activity-dependent manner

Our finding that Hrs puncta exhibit a >2-fold increase in anterograde transport following 2h Bic/4AP treatment (Fig. 2F) implicates a plus-end directed kinesin motor in the activity-dependent transport of Hrs. Previous studies identified several kinesins responsible for the anterograde transport of early endosomes, including kinesin-3 family members KIF13A and KIF13B (Bentley et al. 2015; Huckaba et al. 2011). Using an assay developed to evaluate the ability of these and other kinesins to transport various cargo (Bentley and Banker 2015), we examined whether KIF13A and/or KIF13B mediate Hrs transport (Fig. 4A). We first confirmed the efficacy of this assay in Neuro2a (N2a) cells, by transfecting cells with the minus-end directed motor dynein-binding domain of BicD2 (BicD2-GFP-FKBP), together with the cargo binding domain of KIF13A or KIF13B (KIF13A-FRB-Myc, KIF13B-FRB-Myc). As previously reported, addition of membrane-permeant rapamycin analog to link the FRB and FKBP domains led to the cotransport of BicD2 and KIF13A/B to the cell centrosome (Fig. S7A-B). In addition, we were able to recapitulate transport of Rab5 by both KIF13A and KIF13B as previously reported (Fig. S7C-F; (Bentley et al. 2015)). We next tested the ability of KIF13A and KIF13B to facilitate transport of RFP-Hrs. Surprisingly, while Hrs was readily transported to the centrosome in the presence of linker and KIF13A (Fig. 4B-C), it was not transported in the presence of linker and KIF13B (Fig. 4D-E). These findings indicate a specificity of the transport mechanism for Hrs+ vesicles that does not exist for Rab5+ vesicles. We further tested the specificity of interactions between Hrs and KIF13A/B by coimmunoprecipitation in N2a cells cotransfected with RFP-Hrs or mCherry control, together with the KIF13A- or KIF13B-FRB-Myc constructs. Consistent with our transport assays, we found that Hrs precipitates KIF13A more efficiently than KIF13B (Fig. 4F-G).

**Figure 4.**
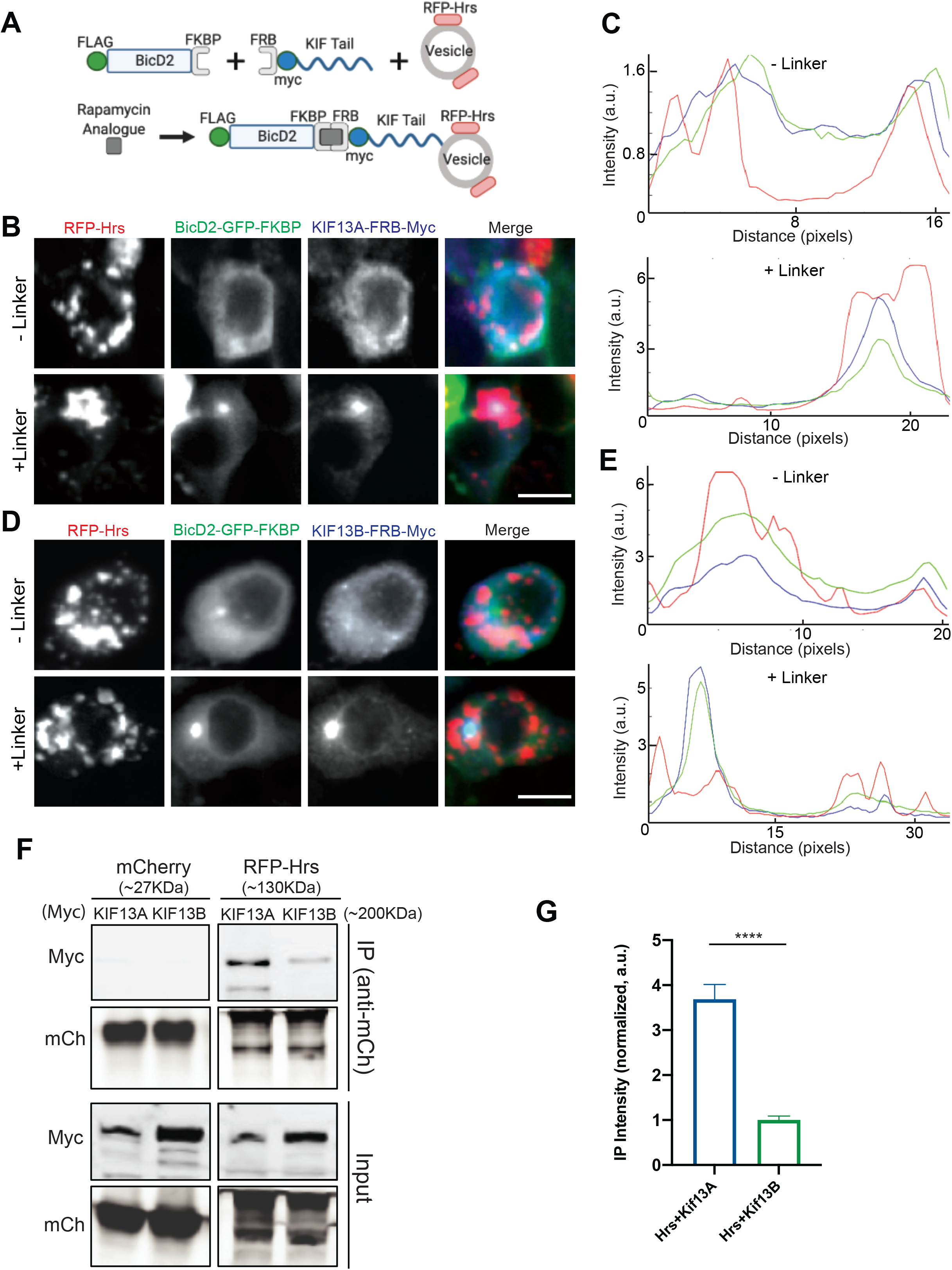
Hrs is transported by KIF13A but not KIF13B. **(A)** Schematic diagram of transport assay utilizing FKBP-tagged BicD2 motor domain, FRB- tagged Kinesin cargo-binding tail domain, and fluorescently-tagged cargo of interest. Upon addition of the rapamycin analogue, FKBP and FRB interact and are co-transported to the centriole. If the protein/vesicle of interest interacts with the Kinesin tail, it is also transported. **(B)** Representative images of N2a cells expressing RFP-Hrs (red), BicD2-GFP-FKBP (green), and KIF13A-FRB-Myc (blue), +/- rapamycin analogue to induce FKBP/FRB interaction (linker). Size bar, 10 µm. **(C)** Line scan graphs of the images in panel **A**, showing that addition of the linker induces colocalization of Hrs with BicD2 and KIF13A as depicted by a single peak in the lower graph. **(D-E)** Representative images of N2a cells expressing RFP-Hrs (red), BicD2-GFP-FKBP (green), and KIF13B-FRB-Myc (blue), +/- linker (**D**), and corresponding line scan graphs (**E**) showing that addition of linker does not induce Hrs colocalization with BicD2/KIF13B. **(F)** Immunoblot of lysates from N2a cells coexpressing mCherry or RFP-Hrs together with KIF13A- FRB-Myc or KIF13B-FRB-Myc, immunoprecipitated (IP) with mCherry antibody and probed with Myc or mCherry antibodies. **(G)** Quantitative analysis of co-immunoprecipitated KIF13A/B, normalized to immunoprecipitated RFP-Hrs and expressed as intensity normalized to KIF13B (****P<0.0001, unpaired t-test, n= 5 independent experiments). Bars show mean ± SEM.

Since anterograde transport of Hrs is regulated by neuronal activity, we next asked whether binding of Hrs to KIF13A is similarly regulated. First, we used the proximity ligation assay (PLA) to visualize interactions between EGFP-Hrs and endogenous KIF13A in the soma of neurons under control or 2h Bic/4AP conditions. In EGFP-expressing neurons, few PLA puncta were present in the control condition, and neuronal activity did not change this value (Fig. 5A, C), indicating little non-specific EGFP/KIF13A interaction under either condition. In contrast, EGFP- Hrs-expressing neurons exhibited a significant increase in PLA puncta following 2h Bic/4AP treatment (Fig. 5B-C), suggestive of an activity-dependent interaction between Hrs and KIF13A. To further confirm the activity-dependence of this interaction, we performed co-immunoprecipitation experiments in N2a cells expressing RFP-Hrs and KIF13A-FRB-Myc following treatment with control or high K+ Tyrodes solution to mimic neuronal depolarization. Consistent with the PLA results, more KIF13A was co-immunoprecipitated with Hrs in cells treated with high K+ Tyrodes (Fig. 5D-E). Together, these findings identify KIF13A as a candidate for the activity-dependent transport of Hrs.

**Figure 5.**
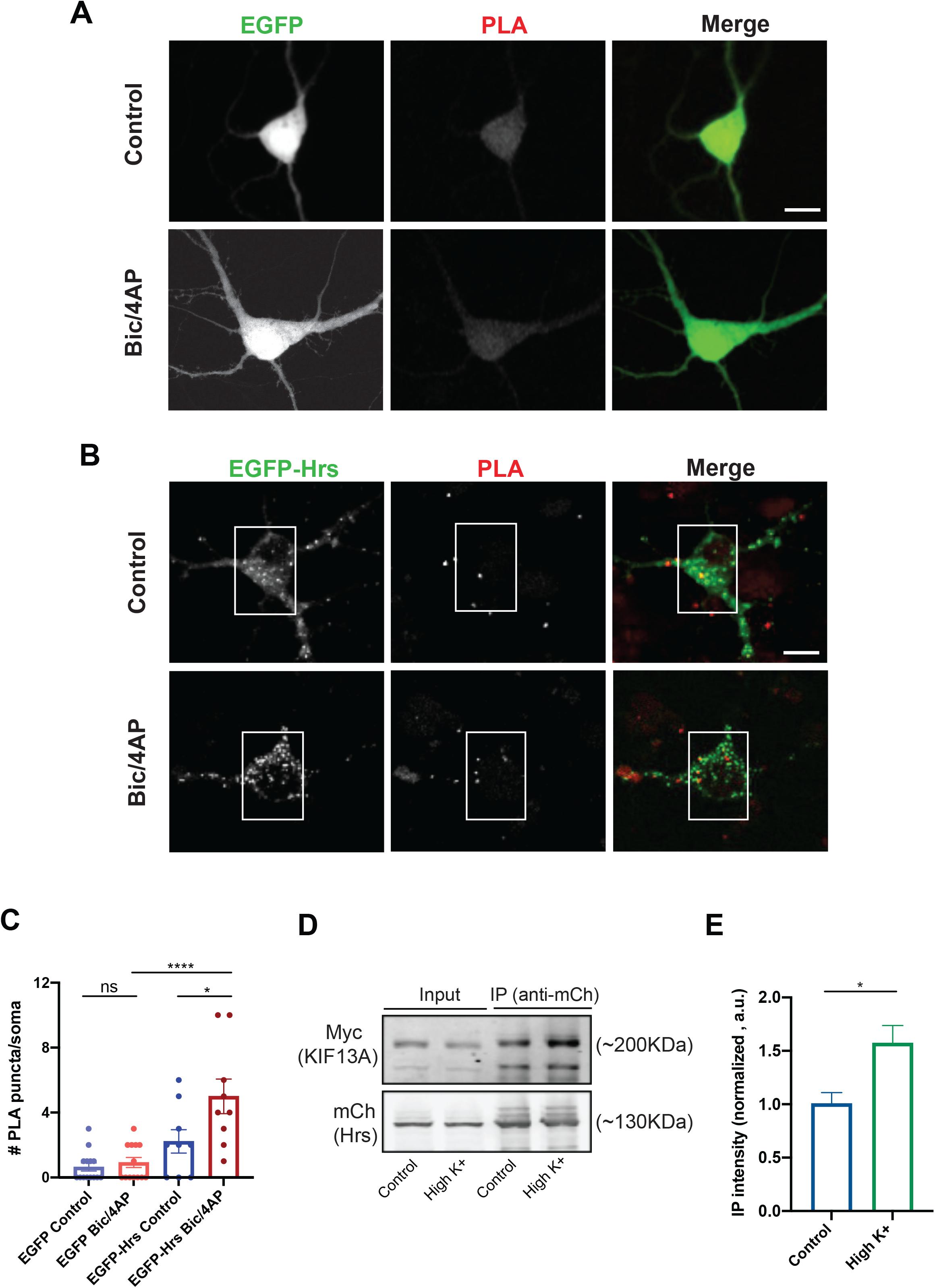
Activity induces the interaction of Hrs with KIF13A. **(A-B)** Representative images for proximity ligation assay (PLA) in 13-15 DIV hippocampal neurons expressing EGFP (**A**) or EGFP-Hrs (**B**) under control or 2hrs Bic/4AP conditions. PLA puncta (red), representing EGFP-Hrs/KIF13A interactions, were quantified in the soma (white rectangles). Size bar, 10 μm. **(C)** PLA puncta number per soma, showing a significant increase in Hrs/KIF13A interactions following 2hrs Bic/4AP treatment (*P=0.0129, ****P<0.0001, one-way ANOVA with Sidak’s multiple comparisons test; n=13 for each EGFP control conditions, n=9 for each EGFP-Hrs conditions, ≥3 independent experiments). **(D)** Immunoblot of lysates from N2a cells coexpressing RFP-Hrs and KIF13A-FRB-Myc, treated with control or high K+ Tyrodes solution for 2hrs, immunoprecipitated (IP) with mCherry antibody and probed with Myc or mCherry antibodies. **(E)** Quantitative analysis of co-immunoprecipitated KIF13A, normalized to immunoprecipitated RFP-Hrs and expressed as intensity normalized to control condition (*P=0.0249, unpaired t-test, 4 independent experiments). Bars show mean ± SEM.

### Knockdown of KIF13A inhibits activity-dependent Hrs motility

To test whether KIF13A is necessary for the axonal transport of Hrs, we examined the effect of KIF13A knockdown on Hrs motility in axons of 13-15 DIV neurons. For these studies, we utilized shRNAs against rat KIF13A (shKIF13A1), which led to ∼50% knockdown of the protein when expressed from 2-15 DIV via lentivirus (Fig. S8A-B). We first examined the effects of KIF13A knockdown on Hrs motility, by coexpressing EGFP-Hrs together with shKIF13A1 or a control shRNA (shCtrl) in neurons plated in microfluidic chambers. Time-lapse imaging revealed that expression of shKIF13A1 did not alter the motility (Fig. 6A-C) or speed (data not shown) of Hrs puncta under control conditions compared to shCtrl. However, shKIF13A1 completely abolished the activity-dependent increase in Hrs motility (Fig. 6A-C, Videos 8-11). We further verified this effect with a second KIF13A hairpin (shKIF13A2; Fig. 6D, S8A-B, E). In contrast, we saw no effect of KIF13B knockdown (by shKIF13B; Fig. S8C-D) on the basal or activity-dependent motility of Hrs (Fig. 6D, S8C-E). Overall, these data indicate that KIF13A, but not KIF13B, is responsible for the activity-dependent transport of Hrs+ vesicles in axons.

**Figure 6.**
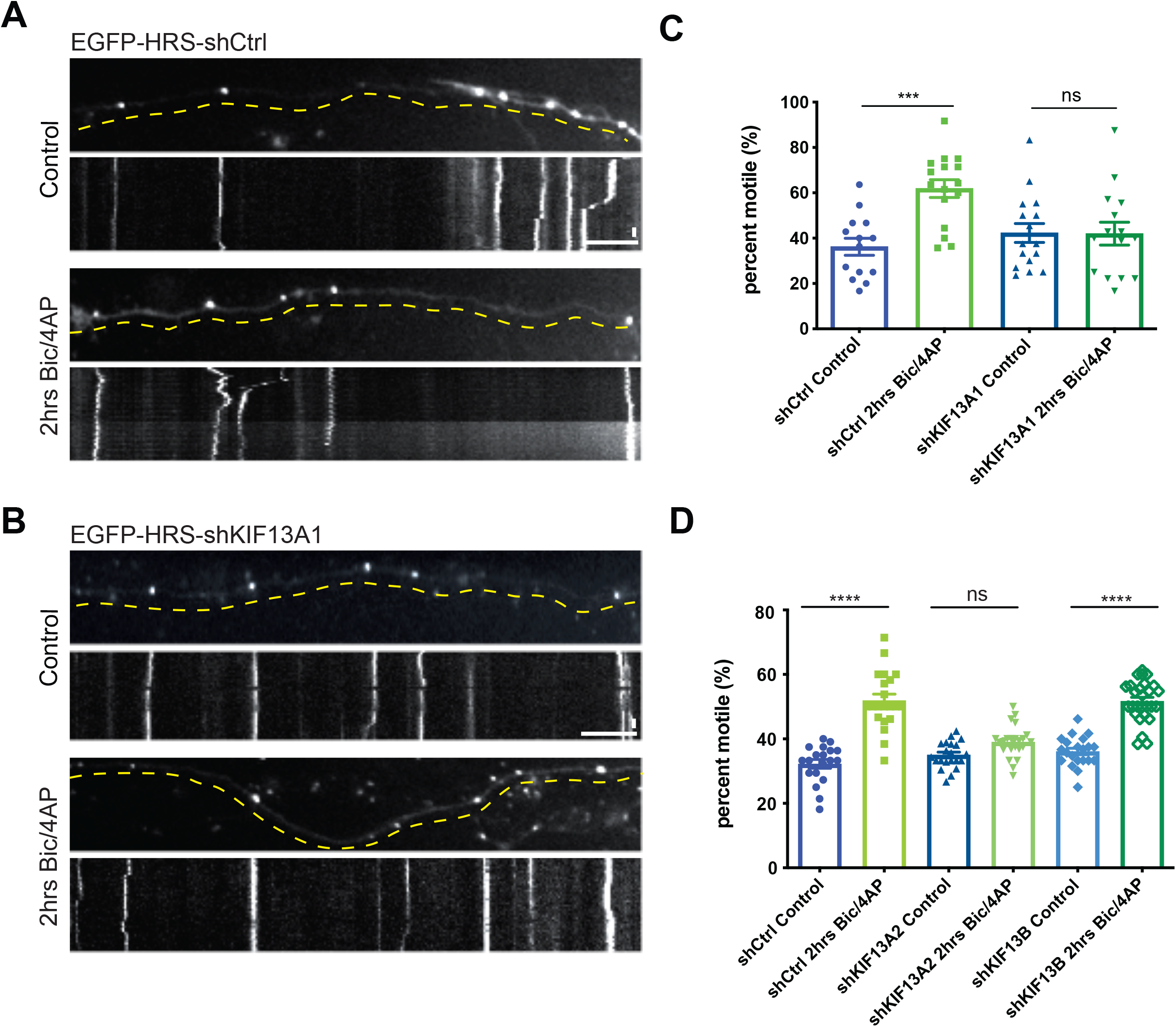
Knockdown of KIF13A inhibits activity-dependent motility of Hrs. **(A)** Representative images and corresponding kymographs of EGFP-Hrs in microfluidically isolated axons expressing shCtrl under control conditions (upper panels) and after 2hrs treatment with Bic/4AP (lower panels)(see Video 8&9). **(B)** Representative images and corresponding kymographs of EGFP-Hrs in microfluidically isolated axons expressing shKIF13A1 under control conditions (upper panels) and after 2hrs treatment with Bic/4AP (lower panels)(see Video 10&11). Horizontal size bar =10 µm, vertical size bar =25 s. 1 frame taken every 5 s (0.2 fps). Dashed yellow lines highlight axons. **(C)** Percentage of motile Hrs puncta in axons, showing that shKIF13A1 prevents the increase in motility induced by 2hrs Bic/4AP treatment (***P=0.0004, ns P=0.9985, one-way ANOVA with Sidak’s multiple comparisons test; ≥3 separate experiments, n=14 (shCtrl control), 16 (shCtrl 2hrs Bic/4AP), 16 (shKIF13A1 control), 15 (shKIF13A1 2hrs Bic/4AP) videos/condition). **(D)** Percentage of motile Hrs puncta in axons expressing shCtrl, shKIF13A2 or shKIF13B under control or 2hrs Bic/4AP treatment. Expression of shKIF13A2, but not shKIF13B, prevents the increase in motility induced by 2hrs Bic/4AP treatment (****P<0.0001, ns P=0.0823, one-way ANOVA with Sidak’s multiple comparisons test; 3 separate experiments, n≥20 videos/condition). Bars show mean ± SEM.

We next investigated whether KIF13A is required for the activity-dependent delivery of Hrs to SV pools. Here, we monitored the colocalization of mCh-Hrs with EGFP-Synapsin in neurons transfected with shCtrl, shKIF13A1, or shKIF13A2 on 5 DIV and treated with vehicle control or Bic/4AP for 20 hrs on 13 DIV. In shCtrl neurons, Bic/4AP treatment significantly increased the colocalization of Hrs with Synapsin (Fig. 7A, D), as anticipated based upon our previous findings (Sheehan et al. 2016). However, no such increase in colocalization was observed in neurons expressing shKIF13A1 or shKIF13A2 (Fig. 7B-D), demonstrating that KIF13A is essential for the activity-induced recruitment of Hrs to SV pools. To determine whether this disruption of Hrs transport has an effect on SV protein degradation, we used our previously-described cycloheximide-chase assay (Sheehan et al. 2016) to measure the degradation of SV membrane proteins Synaptic Vesicle protein 2 (SV2) and Vesicle-associated membrane protein 2 (VAMP2) in neurons expressing shKIF13A1 vs shCtrl. Indeed, the degradation of both SV proteins was slowed in the presence of shKIF13A1 (Fig. 7E-G), suggesting that KIF13A is necessary for facilitating degradation of these SV membrane proteins by delivering Hrs to presynaptic terminals.

**Figure 7.**
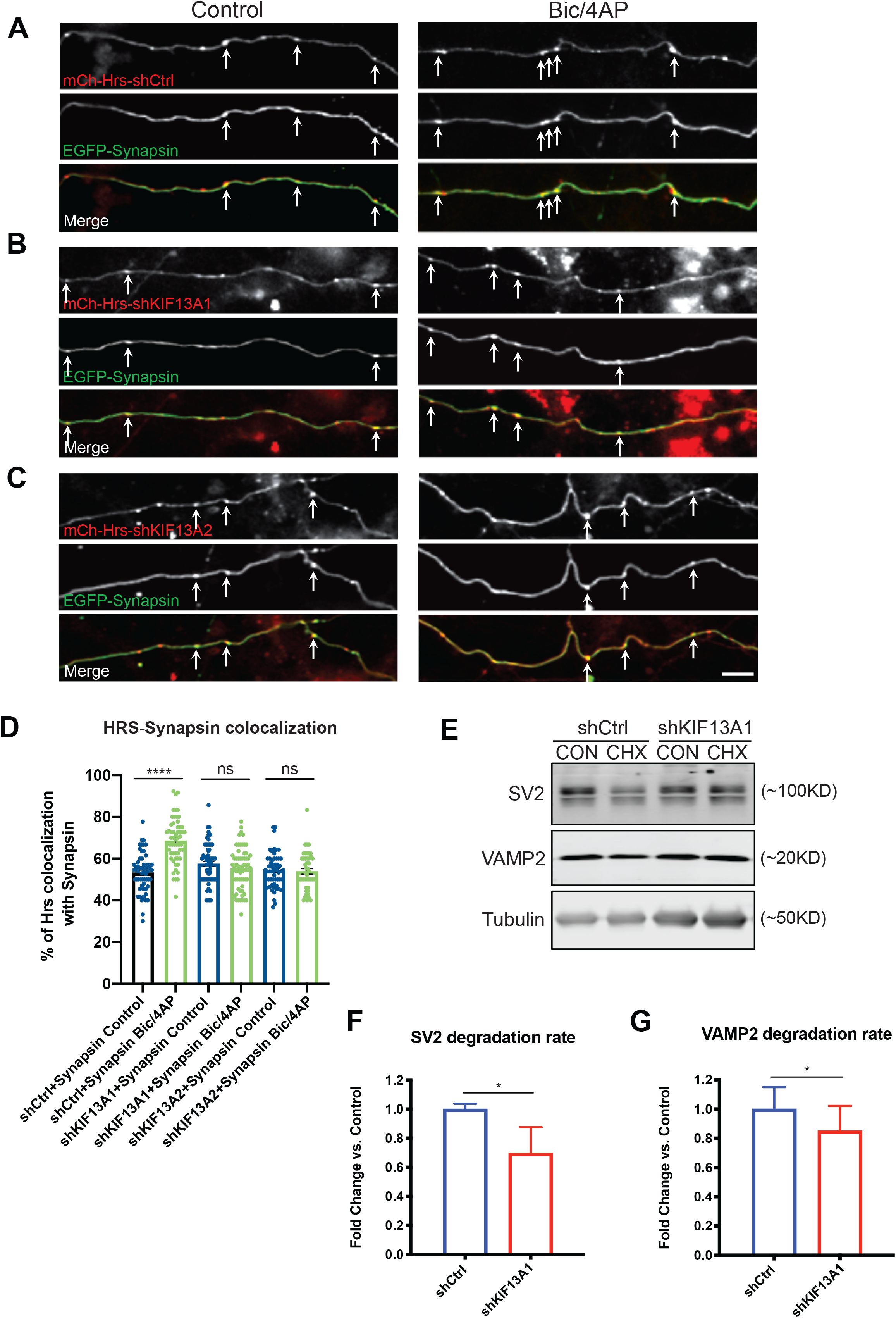
Knockdown of KIF13A impairs activity-dependent transport of Hrs to SV pools. **(A-C)** Representative images of axons expressing mCh-Hrs/shCtrl (**A**), mCh-Hrs/shKIF13A1 (**B**), or mCh-Hrs/shKIF13A2 (**C**) together with EGFP-Synapsin, under control conditions (left panels) or after 20hrs treatment with Bic/4AP (right panels). Size bar, 10 µm. **(D)** Percentage of Hrs puncta colocalized with Synapsin under the conditions shown in A-C, demonstrating that KIF13A knockdown prevents activity-dependent recruitment of Hrs to SV pools (****P<0.0001, ns P=0.0823, one-way ANOVA with Sidak’s multiple comparisons test; 3 separate experiments, n≥20 videos/condition). **(E)** Representative immunoblots of SV2 and VAMP2, with corresponding tubulin loading controls, from lysates of 14 DIV hippocampal neurons expressing shCtrl or shKIF13A1 and treated for 24hrs with DMSO control (CON) or cycloheximide (CHX). **(F-G)** Quantification of the fold-change in SV2 (**F**) and VAMP2 (**G**) degradation under these conditions, calculated as previously described (Sheehan et al. 2016). Degradation of both proteins is significantly decreased in the presence of shKIF13A1 (*P=0.0191 (SV2), *P=0.0161 (VAMP2), unpaired t-test; ≥3 separate experiments, n = 5 (SV2), 11 (VAMP2) cell lysates/condition). All scatter plots show mean ± SEM.

## Discussion

In this study, we demonstrate a novel mechanism for spatiotemporal regulation of the ESCRT pathway in neurons. Our findings show that Hrs transport vesicles exhibit increased motility in axons and delivery to synapses in response to neuronal firing (Fig. 8), providing the first example of activity-dependent transport of degradative machinery in axons. Previous studies have reported that the axonal motility of mitochondria and the inter-bouton transport of SVs in hippocampal neurons are regulated by neuronal activity, specifically presynaptic calcium influx (Lin and Sheng 2015; Chen and Sheng 2013; Gramlich and Klyachko 2017; Qu et al. 2019). These processes are hypothesized to contribute to the activity-dependent remodeling of synapses to alter neurotransmitter release properties (Lin and Sheng 2015; Gramlich and Klyachko 2017). Synaptic activity has also been shown to induce the postsynaptic recruitment of proteasomes and lysosomes in cultured neurons, as well as the local biogenesis of autophagosomes at pre- and postsynaptic sites (Wang et al. 2015; Bingol and Schuman 2006; Goo et al. 2017; Shehata et al. 2012; Soukup et al. 2016; Vanhauwaert et al. 2017; Kulkarni and Maday 2018). However, the axonal transport of autophagosomes in cultured neurons appears to be a largely constitutive process that is unaffected by stimuli such as synaptic activity and glucose levels (Maday and Holzbaur 2016; Maday, Wallace, and Holzbaur 2012). Axonal autophagosomes, once formed, undergo reliable retrograde trafficking to the soma with their cargo while acidifying into autolysosomes (Maday and Holzbaur 2016; Maday, Wallace, and Holzbaur 2012). In contrast, we find that axonal transport of ESCRT-0 proteins Hrs and STAM1 is bidirectional and highly sensitive to neuronal activity levels. These characteristics likely facilitate increased contact of this early ESCRT machinery with SV pools, thereby catalyzing the activity-dependent degradation of SV proteins.

**Figure 8.**
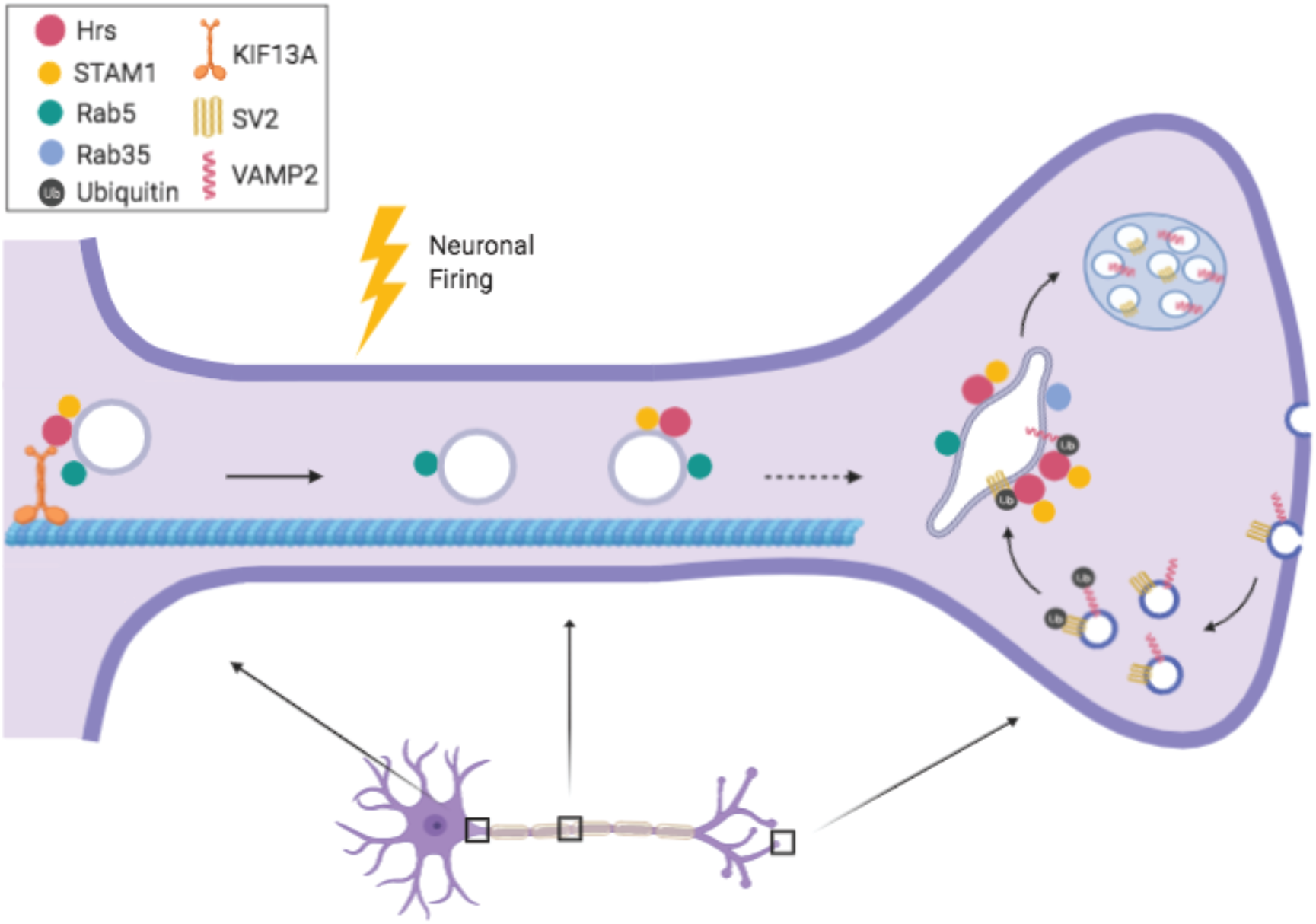
Model for how neuronal activity stimulates the transport of Hrs to SV pools. Neuronal firing increases the association of Hrs with plus-end directed kinesin motor protein KIF13A, stimulating the anterograde transport of these vesicles and their delivery to SV pools. Hrs+ vesicles also transport STAM1 and are a subset of Rab5+ early endosomes.

Our data suggest that axonal Hrs+ vesicles represent a specialized endosomal subtype. Endosomes are highly dynamic and heterogenous structures in neurons, whose complex morphology and high protein trafficking demands have given rise to multiple distinct endosomal classes that are spatially restricted within different neuronal compartments (Maday and Holzbaur 2016; Yap et al. 2018). Early endosomes are classically defined by the markers Rab5 and EEA1 (Langemeyer, Fröhlich, and Ungermann 2018; Simonsen et al. 1998; Gorvel et al. 1991); however, recent studies have identified sub-categories of early endosomes in neurons that can be differentiated both by their markers and axonal transport patterns (Olenick, Dominguez, and Holzbaur 2018; Leonard et al. 2008). The Hrs vesicles that we describe here represent a novel addition to this category. We find that while the total pool of axonal Rab5+ vesicles exhibits high motility under baseline conditions and relatively modest activity-induced increases in anterograde motility, the Hrs+ subset of this pool exhibits limited motility at baseline, but significant activity-induced anterograde and bidirectional motility (Figs. 2 and 3). Moreover, our CLEM experiments demonstrate that Hrs is associated with degradative structures (Fig. 1), though we cannot yet correlate the morphology of these vesicles with their axonal transport history to determine what stage of the degradative process they represent. While we did not assess STAM1 vesicle morphology or colocalization with Rab5 in this study, we did observe that co-expressed Hrs and STAM1 are highly colocalized in axons (Fig. 2), and that STAM+ vesicles exhibit very similar activity-dependent dynamics to Hrs+ vesicles (Fig. S4). This unique sensitivity of ESCRT-0 vesicles to neuronal activity, together with their activity-induced transport to SV pools, suggests that they have a specific function: mediating the activity-dependent degradation of SV proteins. This hypothesis, to be further explored in future studies, aligns with previous research indicating that SV proteins become increasingly dysfunctional as they undergo multiple cycles of exo/endocytosis (Truckenbrodt et al. 2018), necessitating their targeting for degradation.

In addition to characterizing the dynamic behavior of Hrs+ transport vesicles, this study investigates the mechanism of Hrs targeting to these vesicles. As in non-neuronal cells (Gaullier et al. 2000; Komada and Soriano 1999), we find that Hrs association with vesicles is dependent upon interactions between its FYVE domain and the endosomal lipid PI3P (Fig. S5). Blocking PI3P synthesis with SAR405 or expressing the Hrs FYVE domain-inactivating mutant R183A both lead to a significant reduction of axonal Hrs vesicles (Fig. S5). However, these manipulations do not completely abolish Hrs association with vesicles in axons. Moreover, the FYVE domain alone does not recapitulate Hrs axonal transport dynamics, including its activity-dependent motility (Fig. S5). Together, these findings indicate that other domain(s) of Hrs mediate its vesicle association and activity-dependent transport. One possibility is the C-terminal coiled-coil domain, required for Hrs membrane targeting in non-neuronal cells (Raiborg et al. 2001). It is interesting to note that EEA1, which targets to early endosome membranes via its FYVE domain, does not localize to axons (Gaullier et al. 2000; Lawe et al. 2000; Wilson et al. 2000), indicating that additional domains and protein interactions may be required for axonal targeting of FYVE domain-containing proteins.

Finally, our work has elucidated the mechanism of activity-dependent transport of Hrs+ vesicles, in particular the role of kinesin-3 family member KIF13A in this process. Surprisingly, we find that although both KIF13A and KIF13B mediate transport of Rab5 in a reconstituted transport assay, as previously reported (Fig. S7; Bentley et al., 2015, Journal of Cell Biology), only KIF13A transports Hrs (Fig. 4). This specificity is confirmed by coimmunoprecipitation assays in which Hrs precipitates KIF13A much more robustly than KIF13B, further bolstering the concept that Hrs+ vesicles comprise a specialized subset of pro-degradative endosomes with unique transport properties. KIF13A and KIF13B are highly homologous proteins that share multiple cargoes, including early and recycling endosome associated proteins Rab5, Rab10, and transferrin receptor (Jenkins et al. 2012; Bentley et al. 2015; Etoh and Fukuda 2019). These kinesins also have distinct cargoes, including low density lipoprotein receptor (for KIF13B; and serotonin type 1a receptor (for KIF13A; (Jenkins et al. 2012; Zhou et al. 2013)), as well as Hrs. It is possible that these distinct cargoes reflect differences in the subcellular localizations of KIF13A and KIF13B in neurons. However, current reports about their localization are conflicting, with one study showing that KIF13A localizes to axons and dendrites while KIF13B is primarily dendritic (Jenkins et al. 2012), and another showing that KIF13B has an axonal bias whereas KIF13A is exclusively dendritic (Yang et al. 2019). In our experiments, we find that knockdown of KIF13A, but not KIF13B, prevents the activity-induced motility of Hrs in axons (Fig. 6) as well as its delivery to SV pools (Fig. 7), indicating that KIF13A alone has a role in the axonal transport of Hrs. Our findings also highlight the importance of KIF13A-mediated transport in SV protein degradation, which is impaired by KIF13A knockdown under basal conditions (Fig. 7). These data suggest that activity- dependent transport of Hrs occurs even under conditions of lower neuronal firing, but is likely difficult for us to capture during our 4-min imaging sessions. Additional work is needed to identify the other motor proteins responsible for ESCRT-0 vesicle motility, including the dynein adaptors involved in retrograde transport of these vesicles.

The ESCRT pathway plays a vital role in maintaining neuronal health and is intimately linked to neurodegenerative disease etiology. Here, we have illuminated a novel mechanism for spatiotemporal regulation of the ESCRT pathway in neurons, namely the activity-induced transport of Hrs to presynaptic SV pools by the kinesin motor protein KIF13A. These findings broaden our understanding of how protein degradation is regulated in neurons, and shed light on how disruption of this degradative pathway may contribute to the etiology of neurodegenerative disease.

## Materials and methods

### Cell Culture

Hippocampal neurons were prepared from E18 Sprague Dawley rat embryos of both sexes using a modified Banker culture protocol (Kaech & Banker, 2006, Nature Protocols; Sheehan et al., 2016, The Journal of Neuroscience; Waites et al., 2009, The Journal of Neuroscience). Neurons were dissociated in TrypLE Express (ThermoFisher/Invitrogen) for 20min, washed with Hanks Balanced Salt Solution (Sigma), and plated in Neurobasal medium (ThermoFisher/Invitrogen) with N21 supplement (R&D Systems) and Glutamax (ThermoFisher/Invitrogen) at a density of 250,000 neurons per well (12 well plates), coverslip (22 x 22 mm square) or 35mm glass-bottom dish (MatTek), or at a density of 100,000 neurons per microfluidic chamber. Coverslips and dishes were pre-coated with 0.25 mg/mL poly-L-lysine (Sigma-Aldrich) in 100 mM boric acid (pH 8.0) for 24 h before plating. Neuro2a (N2a) neuroblastoma cells (ATCC CCL-131) and HEK293T cells (Sigma) were cultured in DMEM-GlutaMAX (ThermoFisher/Invitrogen) with 10% FBS (Atlanta Biological) and Antibiotic-Antimycotic (ThermoFisher/Invitrogen) and kept at 37°C in 5% CO2.

### Microfluidic Chambers

Microfluidic molds and chambers were prepared as described in (Birdsall et al., 2019). Chambers were prepared by pouring ∼30mL of PDMS mixture (QSil 216) into molds, then baking at 70° C overnight.

### DNA Constructs

pVenus-2xFYVE was a gift from F. Polleux and was subcloned into pFUGWm vector (Waites et al. 2013) at MfeI/NheI sites; pFU-RFP-Hrs-W, pFU-STAM-mCh-W, pFU-EGFP/mCh-Rab35, and pFU-EGFP/mCh-Rab5 were described in (Sheehan et al. 2016)); pFU-EGFP-Synapsin-W was described in (Leal-Ortiz et al. 2008); and pBa-EGFP-flag-BicD2594-FKBP (Addgene plasmid #63569; http://n2t.net/addgene:63569; RRID:Addgene_63569), pBa-FRB-3myc-KIF13A tail 361- 1749 (Addgene plasmid #64288; http://n2t.net/addgene:64288; RRID:Addgene_64288), and pBa-FRB-3myc-KIF13B tail 442-1826 (Addgene plasmid #64289; http://n2t.net/addgene:64289; RRID:Addgene_64289) were gifts from Gary Banker and Marvin Bentley (Jenkins et al. 2012; Bentley et al. 2015; Bentley and Banker 2015). Full-length human Hrs (NCBI accession #D84064.1) was synthesized at Genewiz and cloned into pFUGWm vector at BsrGI/XhoI sites to create pFU-EGFP-Hrs-W, and EGFP replaced with Halotag/mCherry at the AgeI/BsrGI sites to create pFU-Halo-Hrs-W/pFU-mCherry-Hrs. To create pFU-EGFP-R183A Hrs-W, an 868 nucleotides fragment of Hrs containing the R183A mutation was synthesized at Genewiz and subcloned into the BsrGI/PshAI sites. shRNA scrambled control and KIF13A knockdown constructs (shCtrl, shKIF13A1 and A2, KIF13B) were purchased from Origene (catalog #TL706020). The KIF13B knockdown construct was a gift from Dr. Richard Vallee (Dept of Pathology & Cell Biology, CUMC). After testing knockdown efficacy, the 29mer or 21mer shRNA (lower-case letters) and surrounding loop sequences (capital letters)(shCtrl: CGGATCGgcactaccagagctaactcagatagtactTCAAGAGagtactatctgagttagctctggtagtgcTTTTT; shKIF13A1:CGGATCGagccagacctctatgatagcaaccatcagTCAAGAGctgatggttgctatcatagaggtctggct TTTTT;shKIF13A2:CGGATCGtacgaggagacactgtccacgctaagataTCAAGAGtatcttagcgtggacagtgtctcctcgtaTTTTT;shKIF13B:GATCCctgtggaagtaattgctgcaaTCAAGAGttgcagcaattacttccacagTTTTT) were then re-synthesized and subcloned into pFUGW U6 vector (Waites et al. 2013) at EcoRI/PacI sites. shRNAs were moved into FUG-Hrs-W vector using MluI/PacI digestion.

### Antibodies

The following antibodies were used: VAMP2 rabbit antibody (Synaptic System #104202), SV2 mouse antibody (Developmental Studies Hybridoma Bank), tubulin mouse antibody (Sigma #t9026), mCherry rabbit polyclonal antibody (Biovision #5993), Myc mouse monoclonal (Santa Cruz 9E10), Hrs rabbit antibody (Cell Signaling D7T5N), GFP mouse antibody (Roche #11814460001), KIF13A rabbit antibody (Thermo Fisher Scientific PA5-30874), KIF13B mouse antibody (Santa Cruz 6E11), neurofilament mouse antibody (Developmental Studies Hybridoma Bank 2H3).

### Pharmacological Treatments

Pharmacological agents were used in the following concentrations and time courses: cycloheximide (Calbiochem, 0.2 µg/µl, 24hrs), bicuculline (Sigma, 40 µM, 5 min-2hrs or 20hrs), 4- aminopyridine (Tocris Bioscience, 50 µM, 5 min-2hrs or 20hrs), AP5 (Tocris Bioscience, 50 µM, 1hr), CNQX (Tocris Bioscience, 10 µM, 1hr), TTX (Tocris Bioscience, 0.5 µM, 2hrs), PR-619 (LifeSensors, 50 µM, 2hrs), Rapamycin analogue linker drug (AP21967, Clontech, 100 nM, 3hrs), SAR405 (Millipore Sigma, 1 µM, 24hrs). Unless otherwise indicated, all other chemicals were purchased from Sigma.

### Evaluation of Neuronal Health

Cell health in microfluidic chambers was evaluated using NucRed Dead 647 reagent to identify cells with compromised membranes along with membrane-permeant NucBlue™ Live ReadyProbes™ Reagent (both ThermoFisher) to evaluate the total number of cells. Neurons were treated with these reagents for 15 min then immediately imaged as described (see Live Neuron Imaging).

### Lentiviral Transduction and Neuronal Transfection

Lentivirus was produced using HEK293T cells as previously described (Birdsall et al., 2019; Sheehan et al., 2016). Neurons were transduced with 50-150 µL of lentivirus on 7 DIV, and imaged or fixed on 13-15 DIV. Neurons were transfected as described in (Sheehan et al. 2016) using Lipofectamine 2000 (ThermoFisher/Invitrogen) on 9 DIV. Briefly, for each 22x22 mm coverslip or 35 mm dish, 2.5 µL Lipofectamine was incubated with 62.5 µL Neurobasal for 5 min, combined with 2.5 µg DNA diluted into 62.5 µL Neurobasal for 15 min, then added to neurons for 1 h at 37°C. Neurons were transfected in Neurobasal medium containing 50 µM AP5 and 10 µM CNQX, washed 3X in this medium, and then returned to their original medium following transfection. Lentivirus was fluidically isolated in chambers as previously described (Birdsall et al., 2019). Briefly, ∼50% of the media from the wells to be infected was removed and placed in adjacent wells, then virus was infected into the reduced-volume wells to contain it within those compartments (Fig. S2).

### Electron micrograph analysis of endosomal structures

Electron microscopy of hippocampal synapses was performed by Glebov and Burrone as described (Glebov et al. 2016). Presynapses were defined visually by the presence of synaptic vesicle pools. Endosomal structures were defined as single-membraned structures with a diameter >80nm. Multilamellar structures were defined a structure with more than one visible membrane. Synaptic area and endosomal/multilamellar structure size were measured using FIJI.

### Correlative light and electron microscopy

For correlative fluorescence and electron microscopy experiments, mouse hippocampal neurons were cultured on sapphire disks (Technotrade #616-100) that were carbon-coated in a grid pattern (using finder grid masks; Leica microsystems, #16770162) to locate the region of interest by electron microscopy. Following 1mg/mL poly-L-lysine (Sigma #P2636) coating overnight, neurons were seeded at densities of 2.5×10^2^ cell/mm^2^ in 12-well tissue-culture treated plates (Corning #3512) and cultured in NM0 medium (Neurobasal medium (Gibco #21103049) supplemented with 2% B27 plus (Life Technologies # A3582801) and 1% GlutaMAX (Thermo Fischer #35050061). STAM-Halo, HRS-Halo and vGlut1-pHluorin were lentivirally transduced at 6 DIV, and experiments performed at 14-15 DIV. To induce activity, 40 µM bicuculine (Bic, Tocris #0130/50) and 50 µM 4-aminopyridine (4AP, Tocris #940) were added to the culture media at 37°. Following 2 h incubation, half of the media was changed to NM0 with 2 mM Jenelia-Halo-549 nm (Lavis Lab #SB-Jenelia-Halo-549) and incubated for an additional 30 min at 37°. Neurons on the sapphire disks were washed 3X with PBS, fixed with 4% paraformaldehyde (EMS #15714) and 4% sucrose (Sigma #S0389) in PBS for 20 min at room temperature, washed again 3X with PBS, and mounted on a confocal scanning microscope (Carl Zeiss; LSM880) using a chamber (ALA science #MS- 508S). Z-series of fluorescence images were acquired using a 40x objective lens at 2048x2048 pixel resolution. Differential interface contrast images of the carbon grid pattern and neurons on the sapphire disks were also acquired to locate the cells in later steps.

Following fluorescence imaging, neurons were prepared for electron microscopy. Here, sapphire disks were placed into 12-well plate containing 1 mL of 2% glutaraldehyde (EMS #16530) and 1 mM CaCl2 (Sigma-Aldrich # 63535) in 0.1 M sodium cacodylate (Sigma-Aldrich #C0250) pH 7.4, and incubated for 1 h on ice. Subsequently, the sapphire disks were washed with 1 mL of chilled 1M cacodylate buffer 3X for 5 min each. Neurons were then postfixed for 1 h on ice with 2 mL of 1% osmium tetroxide (EMS #RT19134) and 1% potassium ferrocyanide (Sigma-Aldrich #P3289) in 0.1 M sodium cacodylate, pH 7.4. After washing 3X with 2 mL of water for 5 min, neurons were stained with 1 mL of 2% uranyl acetate (Polysciences #21447-25) in water for 30 min at room temperature and then briefly washed with 50% ethanol (Pharmco #111000200) before dehydration in a graded ethanol series (50%, 70%, 90% for 5 min each, and 100% for 5 min 3x). After dehydration, sapphire disks were embedded into 100% epon-araldite resin (Ted Pella #18005, 18060, 18022) with the following schedule and conditions: overnight at 4 °C, 6 h at room temperature (solution exchange every 2 h), and 48 h at 60 °C.

For electron microscopy imaging, regions of interest were located based on the bright field images. When a sapphire disk is removed, neurons and carbon-coating remain in the plastic block, which was trimmed to the regions of interest and sectioned with a diamond knife using an ultramicrotome (Leica; UC7). Approximately 40 consecutive sections (80 nm each) were collected onto the pioloform-coated grids and imaged on a Phillips transmission electron microscope (CM120) equipped with a digital camera (AMT; XR80). The fluorescence and electron micrographs were roughly aligned based on size of the pixels and magnification, and the carbon-coated grid patterns. The alignment was slightly adjusted based on the visible morphological features.

### Immunofluorescence Microscopy

Neurons or N2a cells were immunostained as described previously (Leal-Ortiz et al. 2008; Sheehan et al. 2016). Briefly, coverslips were fixed with Lorene’s Fix (60 mM PIPES, 25 mM HEPES, 10 mM EGTA, 2 mM MgCl2, 0.12 M sucrose, 4% formaldehyde) for 15 min, incubated with primary antibodies and Alexa-488, -568, or -647-conjugated secondary antibodies diluted in blocking buffer (2% glycine, 2% BSA, 0.2% gelatin, 50 mM NH4Cl in PBS) for 1 h at room temperature, and washed 4X with PBS between antibody incubations. Coverslips were mounted with DAPI VectaShield (Vector Laboratories) and sealed with clear nail polish. Alternatively, coverslips were mounted with Aqua-Poly/Mount (Polysciences) and dried overnight in the dark. Images were acquired with a 40x objective (Neofluar, NA 1.3) or 63x objective (Neofluar, NA 1.4) on an epifluorescence microscope (Axio Observer Z1, Zeiss) with Colibri LED light source, EMCCD camera (Hamamatsu) and Zen 2012 (blue edition) software. Images were obtained and processed using Zen Blue 2.1 software. Image processing was done using FIJI (Image-J).

### In vitro KIF13A/KIF13B transport assay

N2a cells were transfected at 50-70% confluence with BicD2-GFP-FKBP, KIF13A-FRB-Myc or KIF13B-FRB-Myc, and either mCh, mCh-Rab5, or STAM-mCh, using Lipofectamine 3000 (ThermoFisher/Invitrogen) according to manufacturer’s instructions. After 48h, rapamycin analogue (AP21967) or EtOH vehicle control were added to relevant samples as described in (Bentley and Banker 2015). After 3 h of treatment, cells were fixed and imaged as described in **Immunofluorescence Microscopy**.

### Live Imaging

Neurons were imaged at 12-15 DIV after being transduced on 7 DIV, or transfected at 9 DIV. Imaging was conducted in pH-maintaining Air Media, consisting of 250 mL L15 Media (ThermoFisher/Invitrogen) and 225 mL Hanks Balanced Salt Solution (Sigma), supplemented with 55mM HEPES, 2% B27 (ThermoFisher), 0.5mM GlutaMAX (ThermoFisher), 0.1% Pen/Strep, 0.5% Glucose, and 2.5% BME. Neurons were imaged at 37°C with a 40x oil-immersion objective (Neofluar, NA 1.3) on an epifluorescence microscope (Axio Observer Z1, Zeiss) with Colibri LED light source, EMCCD camera (Hamamatsu) and Zen 2012 (blue edition) software. One frame was taken every 5 seconds, for a total of 50 frames. Images were obtained and processed using Zen Blue 2.1 software. Axons were identified via location and morphological criteria. Image processing was done using FIJI (Image-J).

### Live Imaging Analysis

Motility of axonal puncta was analyzed using FIJI’s “Manual Tracker” plugin. Raw data for the movement of each puncta over each video frame were imported into Matlab and motility, directionality, and speed were calculated using a custom program. Only puncta ≥0.5 microns and <1.5 microns in diameter were analyzed. Directionality of each puncta was based on net displacement over the course of the video (Fig. S3F), and was categorized as follows: ≥4 µm away from the cell body = anterograde, ≥4 µm towards the cell body = retrograde, ≥4 µm total, but <4 µm in one direction = bidirectional, <4 µm = stationary. Axon length was measured in FIJI.

### Proximity ligation assay

Proximity ligation assay (PLA) was performed in hippocampal neurons according to manufacturer’s instructions (Duolink, Sigma). Until the PLA probe incubation step, all manipulations were performed as detailed above for the immunocytochemistry procedure. PLA probes were diluted in blocking solution. The primary antibody pairs used were anti-KIF13A (Rabbit, 3 μg/ml) and anti-GFP (Mouse, 1 μg/ml). All protocol steps were performed at 37°C in a humidified chamber, except for the washing steps. Coverslips were then mounted using Duolink In situ Mounting Media with DAPI.

### Immunoprecipitation and Western Blot

For immunoprecipitation assays, N2a cells were transfected at 50-70% confluence using Lipofectamine 3000 (ThermoFisher/Invitrogen) according to manufacturer’s instructions. Cells were collected 48 h later in lysis buffer (50 mm Tris-Base, 150 mm NaCl, 1% Triton X-100, 0.5% deoxycholic acid) with protease inhibitor mixture (Roche) and clarified by centrifugation at high speed (20,000 rcf). Resulting supernatant was incubated with Dynabeads (ThermoFisher/Invitrogen) coupled to anti-mCherry antibodies (polyclonal; Biovision). Lysates and beads were incubated at 4°C under constant rotation for 2 h. Beads were washed 2–3X with PBS containing 0.05% Triton (PBST) and then once with PBS. Bound proteins were eluted using sample buffer (Bio-Rad) and subject to SDS-PAGE immunoblotting as described following. For all other immunoblotting experiments, neurons were collected directly in 2X SDS sample buffer (Bio-Rad). Samples were subjected to SDS-PAGE, transferred to nitrocellulose membranes, and probed with primary antibody in 5% BSA/PBS + 0.05% Tween 20 overnight at 4°C, followed by DyLight 680 or 800 anti-rabbit, anti-mouse, or anti-goat secondary antibodies (ThermoFisher) for 1 h. Membranes were imaged using an Odyssey Infrared Imager (model 9120, LI-COR Biosciences). Protein intensity was measured using the “Gels” function in FIJI.

### Statistical Analyses

All statistical analyses were performed in GraphPad Prism. Statistics for each experiment are described in the Figure Legends. A D’Agostino-Pearson normality test was used to assess normality. An unpaired Student’s t test or a Mann–Whitney U test was used to compared two datasets, and an ANOVA or a Kruskal-Wallis test was used to compare three or more datasets. For multiple comparisons, Dunnett’s or Dunn’s post-hoc test was used to compare a control mean to other means, while Sidak’s post-hoc test was used to compare pairs of means. Statistical significance is noted as follows (and in Figure Legends): ns, P>0.05; *, P≤0.05; **, P≤0.01; ***, P≤0.001, ****, P≤0.0001. All experiments were repeated with at least 3 independent sets of samples.

### Data Availability

The data that support the findings of this study are available from the corresponding author upon reasonable request.

## Supplemental Material

Supplemental Figures 1-8 and Supplemental Figure legends Videos 1-11

## Acknowledgements

This work was supported by NIH grants 2R01NS080967 to C.W. and S.W., 5T32NS064928 to V.B., and Hirschl Research Scientist Award to C.W. We would like to thank Drs. Oleg Glebov and Juan Burrone (King’s College, London, UK) for generously sharing EM micrographs of synapses from hippocampal neurons treated with TTX and gabazine, Dr. Ulrich Hengst (Columbia University) and Hengst lab members for help with microfluidic chambers, and members of the Waites lab for their scientific input. The authors declare no competing financial interests.

## Author Contributions

V.B., M.Z. and C.W. designed the research, V.B., M.Z., K.K., and Y.I. performed experiments and analyzed data, C.W. and S. W. supervised the research, and V.B., M.Z. and C.W. wrote the manuscript. All authors reviewed the manuscript for accuracy and clarity.

## Competing Interests

The authors declare no conflict of interest.

## Abbreviations

ESCRT: endosomal sorting complex required for transport
Hrs: hepatocyte growth factor-regulated tyrosine kinase substrate
MVB: multivesicular body
STAM1: signal transducing adaptor molecule 1
SV: synaptic vesicle

## Notes

### Competing Interest Statement

The authors have declared no competing interest.

### Summary of Updates

Figures were updated to more clearly show axons and to provide better images of fluorescently-tagged Hrs and Rab puncta in axons. New data further illuminates the mechanism of Hrs transport to presynaptic boutons, showing that neuronal activity stimulates the interaction between Hrs and motor protein KIF13A (Figure 5), and that KIF13A is required for the activity-dependent transport of Hrs to synaptic vesicle pools in axons (Figure 7).

