## Supplemental Figure and Video legends for "Axonal transport of Hrs is activity-dependent and rate limiting for synaptic vesicle protein degradation"

**Supplemental Figures and Videos**

**Figure S1: Neuronal activity stimulates recruitment of endogenous Hrs into axons.**

**(A)** Representative images of axons from dissociated hippocampal neurons, immunostained with Hrs and neurofilament antibodies under control conditions (upper panels) and after 20hrs treatment with Bic/4AP (lower panels). Size bar, 10 μm. **(B)** Hrs puncta intensity under control or Bic/4AP treatment conditions, normalized to the control condition (****P<0.0001, unpaired t-test; 3 independent experiments, n=27 (control), 26 (Bic/4AP) fields of view/condition). **(C)** Number of Hrs puncta/100 μm of axon under control or Bic/4AP treatment conditions (****P<0.0001, unpaired t-test; 3 independent experiments, n=38 (control), 46 (Bic/4AP) fields of view/condition). All scatter plots show mean ± SEM.

**Figure S2: Experimental setup and characterization of cell health, lentiviral transduction efficiency, and neuronal activity effects in microfluidic chambers. (A)** Schematic diagram of microfluidic chambers and method of lentiviral transduction. **(B)** Lentiviral transduction efficiency of neurons in microfluidic chambers. Graph shows a similar percentage of cell bodies transduced for each lentivirus (≥3 independent experiments, n=28 (Hrs), 23 (STAM), 26 (Rab5), 29 (Rab35) fields of view/condition). **(C)** Analysis of neuronal health in chambers. Graph shows percentage of cells exhibiting NucRed Dead fluorescence following treatments, which is similar across conditions (ns P=0.2678, one-way ANOVA with Dunnett's multiple comparisons test; ≥3 independent experiments, n=33 (control), 24 (acute activity), 31 (2hrs activity) videos/condition). **(D)** Percentage of axonal Hrs puncta that exhibit motility under control conditions, 2hrs Bic/4AP treatment, and 2hrs TTX treatment to block neuronal activity. TTX treatment does not alter puncta motility compared to the control condition (***P=0.0002, ns P=0.5517, one-way ANOVA with Dunnett's multiple comparisons test; ≥3 separate experiments, n=18 (control), 15 (2hrs Bic/4AP), 31 (2hrs TTX) videos/condition). All scatter plots show mean ± SEM.

**Figure S3: Neuronal activity shifts the net displacement of motile Hrs puncta in the anterograde direction. (A-C)** Representative images and corresponding kymographs of EGFP-Hrs in axons from 13-15 DIV dissociated hippocampal neurons under control conditions (**A**), acute treatment with Bicuculline/4AP (**B**), or 2hrs treatment with Bicuculline/4AP (**C**). Horizontal size bar, 10 µm, vertical size bar, 25 s. 1 frame taken every 5 s (0.2 fps). Dashed yellow lines highlight axons. **(D)** Percentage of motile Hrs puncta in axons, showing a significant increase with acute and 2hrs Bic/4AP treatment (****P<0.0001, **P=0.0043, one-way ANOVA with Dunnett's multiple comparisons test; 3 independent experiments, n=14 (control), 16 (acute Bic/4AP), 14 (2hrs Bic/4AP) videos/condition). **(E)** Average speed of motile Hrs puncta (µm/sec), showing no change across conditions (one-way ANOVA with Dunnett's multiple comparisons test; ≥3 independent experiments, n=16 videos/condition). **(F)** Examples of Hrs puncta exhibiting retrograde (orange line), stationary (purple line), and anterograde (blue line) motility. Size bar is 4µm. All scatter plots and graphs show mean ± SEM. **(G)** Histogram showing relative frequency of net displacements of tracked Hrs puncta (anterograde, retrograde, and bidirectional). Bins=1µm, negative values indicate retrograde displacement. Line shows Gaussian curve fit to data; ≥3 separate experiments, n=20 videos, 229 tracked puncta (DMSO), 16 videos, 252 tracked puncta (2hrs Bic/4AP). **(H)** Histogram showing total frequency of net displacement of tracked Hrs puncta (anterograde, retrograde, and bidirectional). Bins=1µm, negative values indicate retrograde displacement. Line shows Gaussian curve fit to data. Same 'n' as (G).

**Figure S4: Neuronal activity stimulates the transport of STAM1+ vesicles in axons**.

**(A-C)** Representative images and corresponding kymographs of mCh-STAM1 in microfluidically isolated axons from 13-15 DIV hippocampal neurons under control (**A**), acute Bic/4AP (**B**), and 2hrs Bic/4AP (**C**) conditions. Horizontal size bar, 10 µm, Vertical size bar, 25 s. 1 frame taken every 5 s (0.2 fps). Dashed yellow lines highlight axons. **(D)** Percentage of motile STAM1 puncta in axons, showing a significant increase with acute and 2hrs Bic/4AP treatment (****P<0.0001, *P=0.016, one-way ANOVA with Dunnett's multiple comparisons test; ≥3 independent experiments, n=37 (control), 41 (acute Bic/4AP), 19 (2hrs Bic/4AP) videos/condition). **(E)** Average speed of motile STAM1 puncta (μm/sec), showing no change across conditions (ns P=0.9839. Same ‘n’ values as D). **(F)** Breakdown of directional movement of STAM1 puncta, showing that anterograde, retrograde, and bidirectional movement are all significantly increased following 2hrs Bic/4AP treatment (****P<0.0001, ***P=0.0007, **P=0.0039, *P=0.044 (bidirectional), *P=0.0111 (stationary), ns P=0.0857, one-way ANOVA with Dunnett's multiple comparisons test; same ‘n’ values as D). All scatter plots and graphs show mean ± SEM.

**Figure S5: Hrs association with axonal vesicles requires FYVE domain/PI3P interactions. (A)** Representative images of axons from 13-15 DIV hippocampal neurons expressing EGFP-Hrs under control conditions (left panels) or 24hrs SAR405 treatment (middle panels), or EGFP-Hrs R183A FYVE domain inactivating mutant (right panels), and immunostained with neurofilament antibody (red). Size bar, 10 μm**. (B)** Number of EGFP-Hrs puncta per 100 μm of axon under the indicated conditions (****P<0.0001, one-way ANOVA with Dunnett's multiple comparisons test; ≥3 independent experiments; n=51 (EGFP-Hrs control), 48 (24hrs SAR405 treatment), 54 (EGFP-Hrs R183A) fields of view/condition). **(C)** EGFP-Hrs puncta intensity under the indicated conditions, normalized to EGFP-Hrs control condition (****P<0.0001, one-way ANOVA with Dunnett's multiple comparisons test; 3 independent experiments, n=24 fields of view for all three conditions). **(D)** Representative images and corresponding kymographs for EGFP-2xFYVE domain in microfluidically isolated axons from 13-15 DIV hippocampal neurons under control conditions (upper panels), acute treatment with Bic/4AP (middle panels), or 2hrs treatment with Bic/4AP (lower panels). Horizontal size bar, 10 μm, vertical size bar, 25 s. 1 frame taken every 5 s (0.2 fps). **(E)** Percentage of motile 2xFYVE puncta in axons, showing no significant difference across conditions (ns P=0.1463, one-way ANOVA with Dunnett's multiple comparisons test; ≥3 independent experiments, n=11 (control), 12 (acute Bic/4AP), 10 (2hrs Bic/4AP) videos/condition). **(F)** Average speed of motile 2xFYVE puncta (μm/sec), showing no change across conditions (ns P=0.5533, one-way ANOVA with Dunnett's multiple comparisons test; ≥3 separate experiments, same 'n' values as D. **(G)** Breakdown of directional movement of 2xFYVE puncta, expressed as percentage of total puncta. All scatter plots and graphs show mean ± SEM.

**Figure S6: Neuronal activity does not stimulate Rab35 motility in axons.**

**(A-B)** Representative images and corresponding kymographs for EGFP-Rab35 in microfluidically isolated axons from 13-15 DIV hippocampal neurons under control conditions (**A**) or 2hrs treatment with Bicuculline/4AP (**B**). Horizontal size bar, 10 μm, Vertical size bar, 25 s. 1 frame taken every 5 s (0.2 fps). **(C)** Percentage of motile Rab35 puncta in axons, showing no significant difference across conditions (ns P=0.1099, unpaired t-test; ≥3 independent experiments, n= 20 (control), 20 (2hrs Bic/4AP) videos/condition). **(D)** Average speed of motile Rab35 puncta (μm/sec), showing no change across conditions (ns P=0.3271, unpaired t-test; ≥3 independent experiments, same 'n' values as C). **(E)** Breakdown of directional movement of Rab35 puncta, expressed as percentage of total puncta, showing no change across conditions (ns P=0.1099, unpaired t-test; ≥3 independent experiments, same 'n' values as C). **(F)** Percentage of Hrs puncta that are positive for Rab35, showing increased colocalization following 2hrs Bic/4AP treatment (*P=0.0348, unpaired t-test; ≥3 independent experiments, n=36 (control), 39 (2hrs Bic/4AP) fields of view/condition). **(G)** Representative images for aaxons from 13-15 DIV hippocampal neurons expressing EGFP-Synapsin and mCh-Rab35 under control conditions (left panel), or 2hrs treatment with Bic/4AP (right panel). Arrows indicate sites of colocalization. Size bar, 10 μm. **(H)** Percentage of Rab35 puncta that are positive for Synapsin, demonstrating no significant differences between control and 2hrs Bic/4AP treatment (ns P=0.0968 unpaired t-test; ≥2 independent experiments, n=11 (control), 6 (2hrs Bic/4AP) videos/condition) All scatter plots and graphs show mean ± SEM.

**Figure S7: KIF13A and KIF13B mediate transport of Rab5+ vesicles in N2a cells.**

**(A-B)** Representative images of N2a cells coexpressing BicD2-GFP-FKBP (green) and KIF13A-FRB-Myc (blue; **A**) or KIF13B-FRB-Myc (blue; **B**), +/- rapamycin analogue to induce FKBP/FRB binding (linker). Size bar, 10 µm. **(C-D)** Representative images of N2a cells expressing mCh-Rab5 (red), BicD2-GFP-FKBP (green), and KIF13A-FRB-Myc (blue), +/- linker, and corresponding line scan graphs (**D**) showing that addition of linker induces colocalization of Rab5 with BicD2/KIF13A. **(E-F)** Representative images of N2a cells expressing mCh-Rab5 (red), BicD2-GFP-FKBP (green), and KIF13B-FRB-Myc (blue), +/- linker, and corresponding line scan graphs (**F**) showing that addition of linker induces colocalization of Rab5 with BicD2/KIF13B.

**Figure S8: Knockdown of KIF13A and KIF13B in neurons.**

**(A)** Representative immunoblot of lysates from 13-15 DIV hippocampal neurons expressing shKIF13A1, shKIF13A2, or scrambled shRNA control (shCtrl), probed with antibodies against KIF13A and tubulin. **(B)** Quantification of KIF13A band intensity, normalized to tubulin control, with shCtrl or shKIF13A expression (***P=0.0004 (shKIF13A1), ***P=0.0002 (shKIF13A2), one-way ANOVA with Dunnett's multiple comparisons test; n=4 independent experiments). Bars show mean ± SEM. **(C)** Representative immunoblot of lysates from 13-15 DIV hippocampal neurons expressing shKIF13B or scrambled shRNA control (shCtrl), probed with antibodies against KIF13B and tubulin. **(D)** Quantification of KIF13B band intensity, normalized to tubulin control, with shCtrl or shKIF13B expression (***P=0.0007, unpaired t-test; n=4 independent experiments). Bars show mean ± SEM. **(E)** Representative images and corresponding kymographs of EGFP-Hrs in axons from dissociated hippocampal neurons expressing shCtrl (left panels), shKIF13A2 (middle panels), or shKIF13B (right panels), under control conditions (upper panels) and after

2hrs treatment with Bic/4AP (lower panels). Horizontal size bar, 10 μm, Vertical size bar, 25 s. 1 frame taken every 5 s (0.2 fps). Dashed yellow lines highlight axons. Quantification of Hrs motility under these conditions is shown in Figure 6D.

**Supplemental Videos**

**Video 1.** RFP-Hrs motility in microfluidically isolated axons under control conditions. Soma is to the right. Still image from video is in Fig. 2A. All videos were obtained using time-lapse epifluorescence microscopy with one frame every 5 seconds. Video frame rate is 5 fps.

**Video 2.** RFP-Hrs motility in microfluidically isolated axons after 2hrs Bic/4AP treatment. Soma is to the right. Still image from video is in Fig. 2C.

**Video 3**. EGFP-Hrs (green) and mCh-STAM1 (red) motility in axons of dissociated hippocampal cultures under control conditions. Still images from video are in Fig. 2G.

**Video 4.** EGFP-Rab5 motility in microfluidically-isolated axons under control conditions. Soma to the right. All videos were obtained using time-lapse epifluorescence microscopy with one frame every 5 seconds. Video frame rate is 5 fps. Still image from video is in Fig. 3A.

**Video 5.** EGFP-Rab5 motility in microfluidically-isolated axons after 2hrs Bic/4AP treatment. Soma to the right. Still image from video is in Fig. 3C.

**Video 6.** RFP-Hrs (red) and EGFP-Rab5 (green) motility in axons of dissociated hippocampal cultures under control conditions. Still images from video are in Fig. 3G (upper panel).

**Video 7.** RFP-Hrs (red) and EGFP-Rab5 (green) motility in axons of dissociated hippocampal cultures after 2hrs Bic/4AP treatment. Still images from video are in Fig. 3G (lower panel).

**Video 8.** EGFP-Hrs motility in microfluidically-isolated axons expressing shCtrl under DMSO control treatment. Soma is to the right. All videos were obtained using time-lapse epifluorescence microscopy with one frame every 5 seconds. Video frame rate is 5 fps. Still image from video is in Fig. 6A (upper panels).

**Video 9.** EGFP-Hrs motility in microfluidically-isolated axons expressing shCtrl after 2hrs Bic/4AP treatment. Soma is to the right. Still image from video is in Fig. 6A (lower panels).

**Video 10.** EGFP-Hrs motility in microfluidically-isolated axons expressing shKIF13A1 under DMSO control treatment. Still image from video is in Fig. 6B (upper panels).

**Video 11.** EGFP-Hrs motility in microfluidically-isolated axons expressing shKIF13A1 after 2hrs Bic/4AP treatment. Still image from video is in Fig. 6B (lower panels).

Supplemental Figure 1

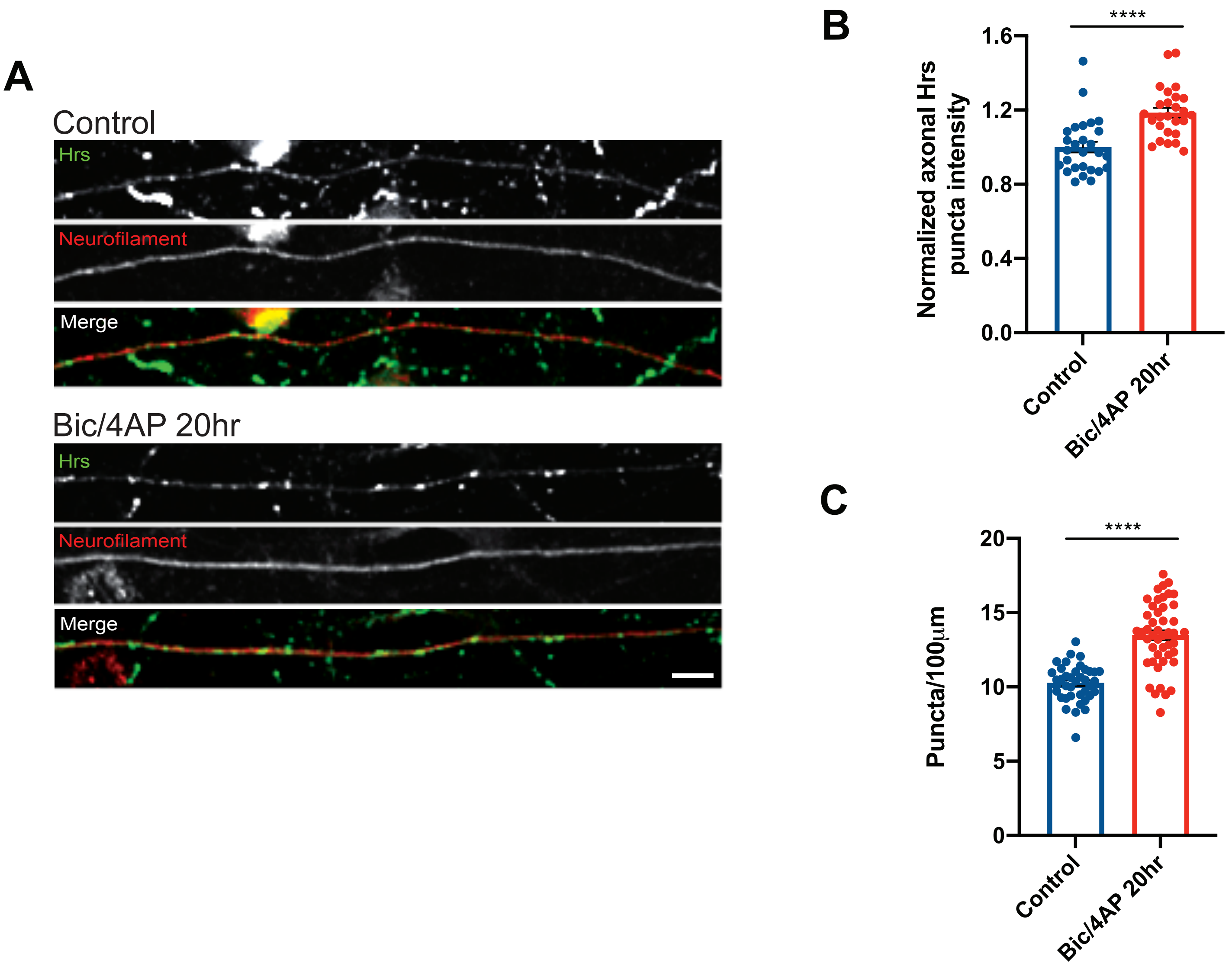

Supplemental Figure 2

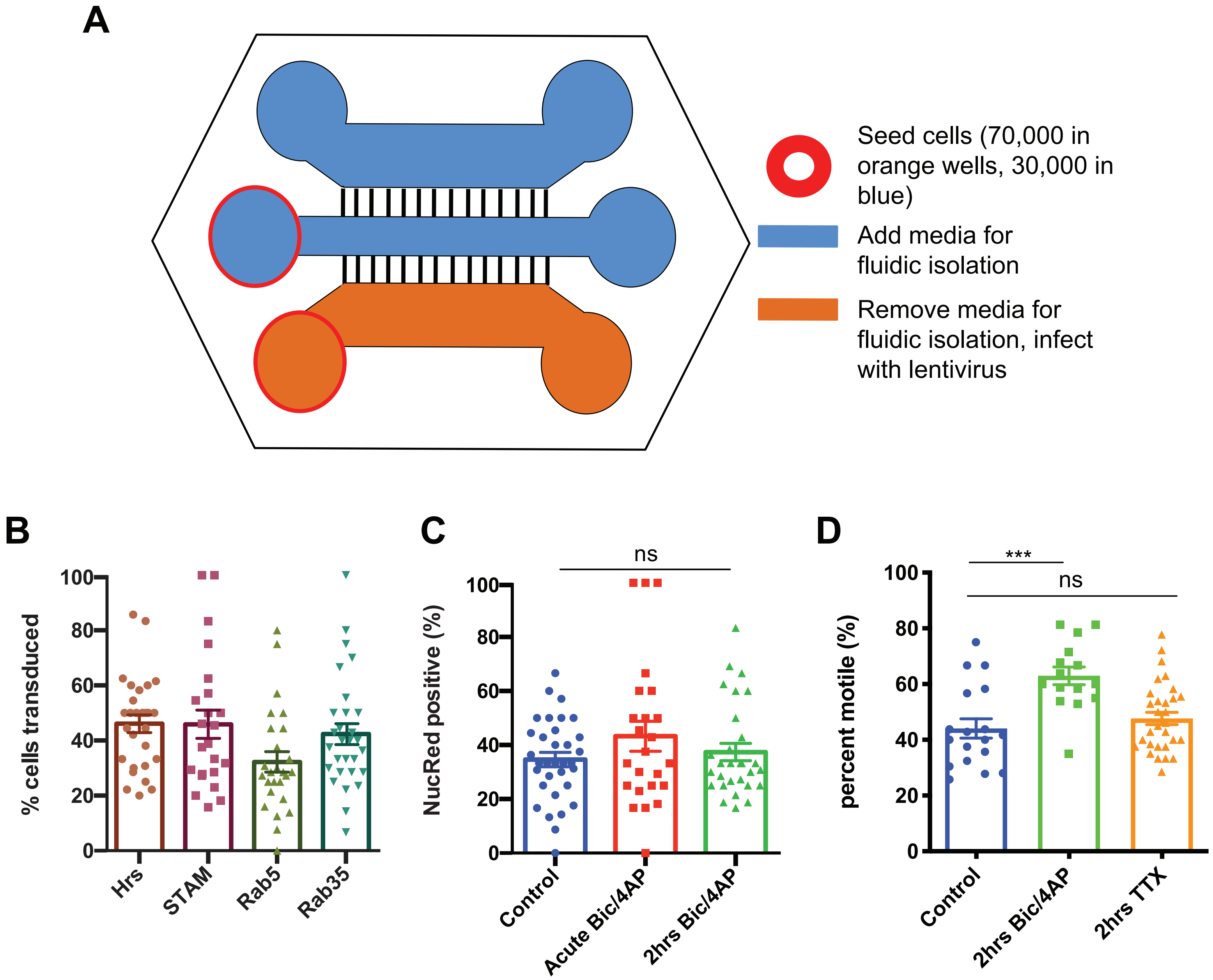

Supplemental Figure 3

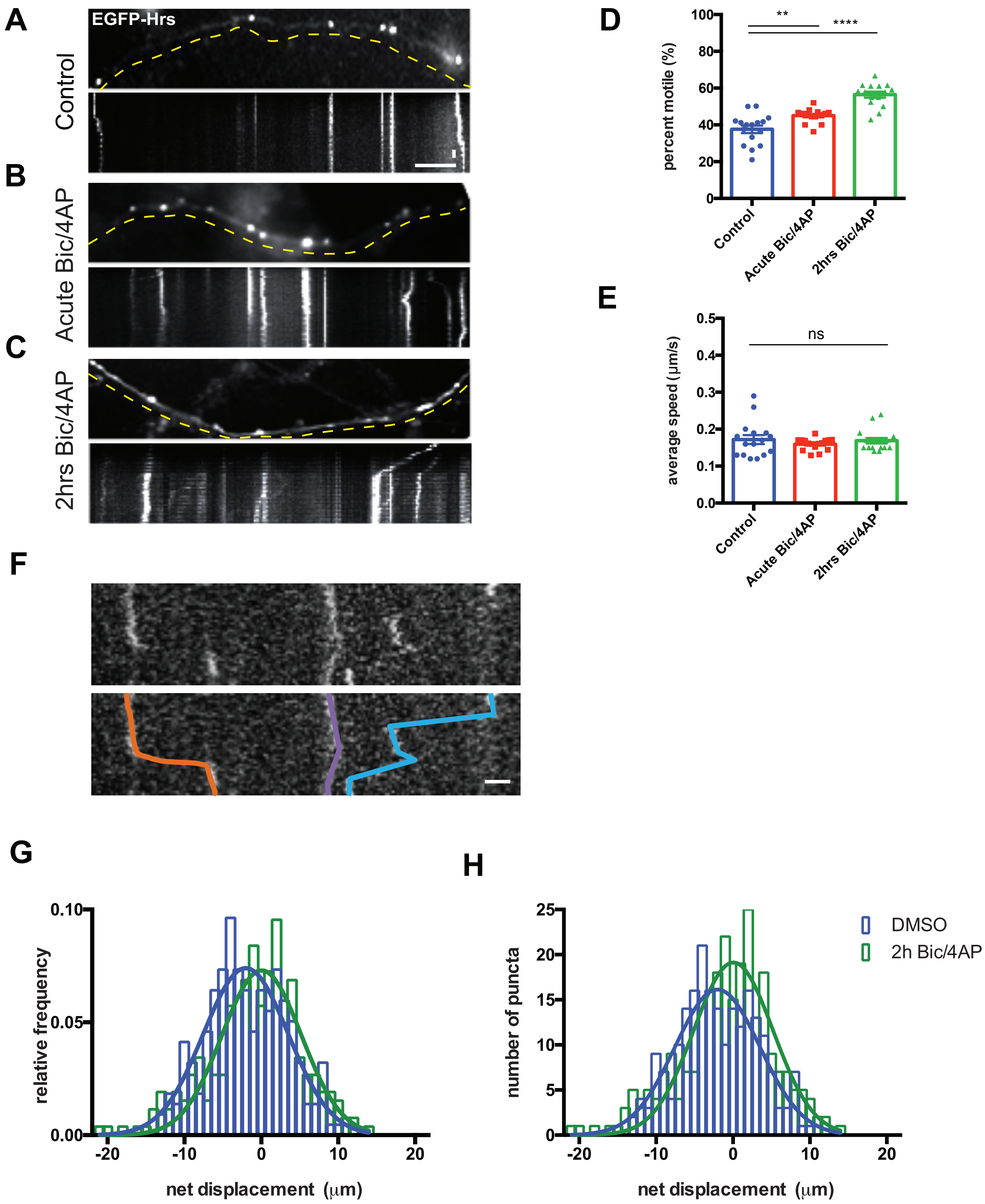

Supplemental Figure 4

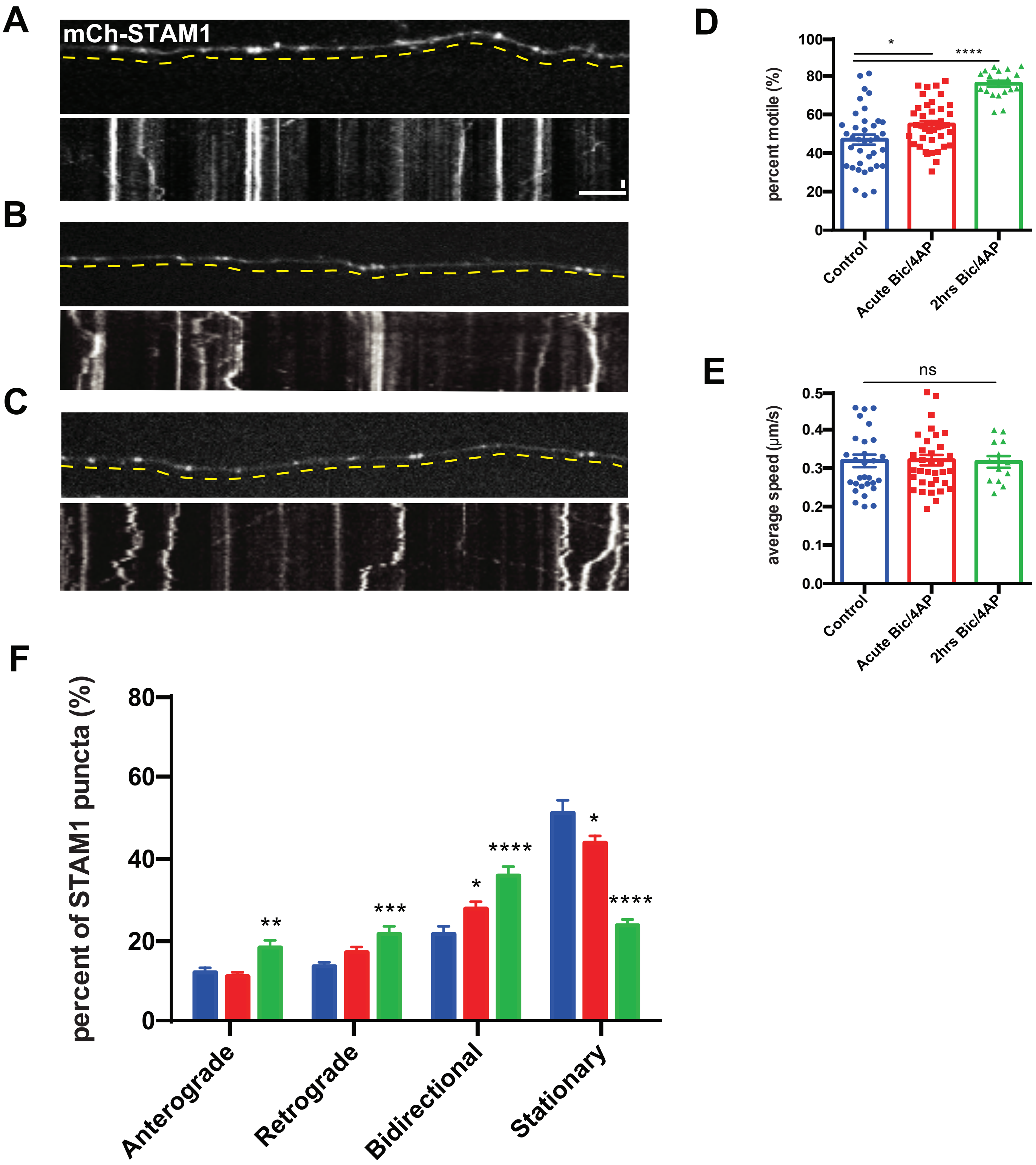

Supplemental Figure 5

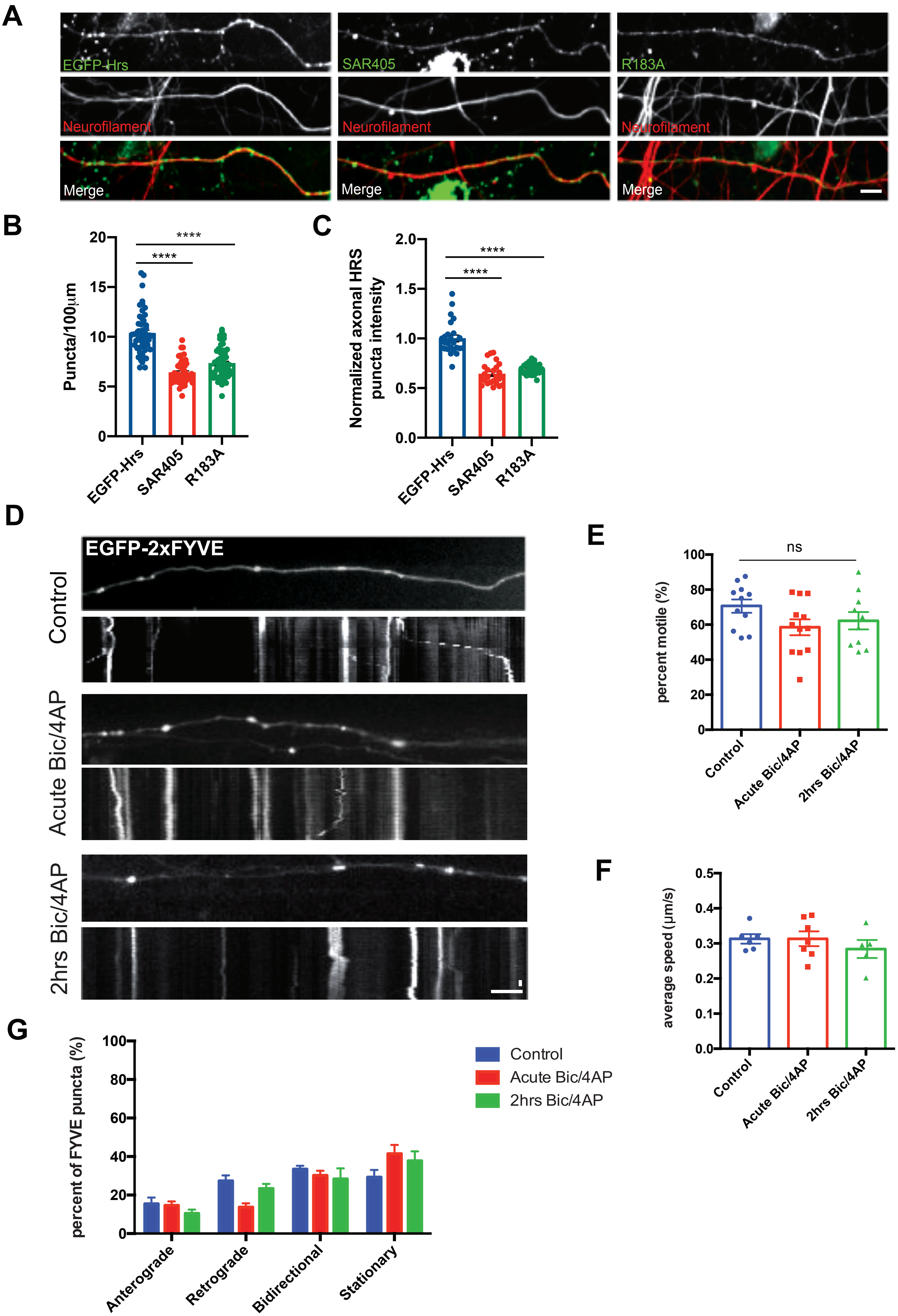

Supplemental Figure 6

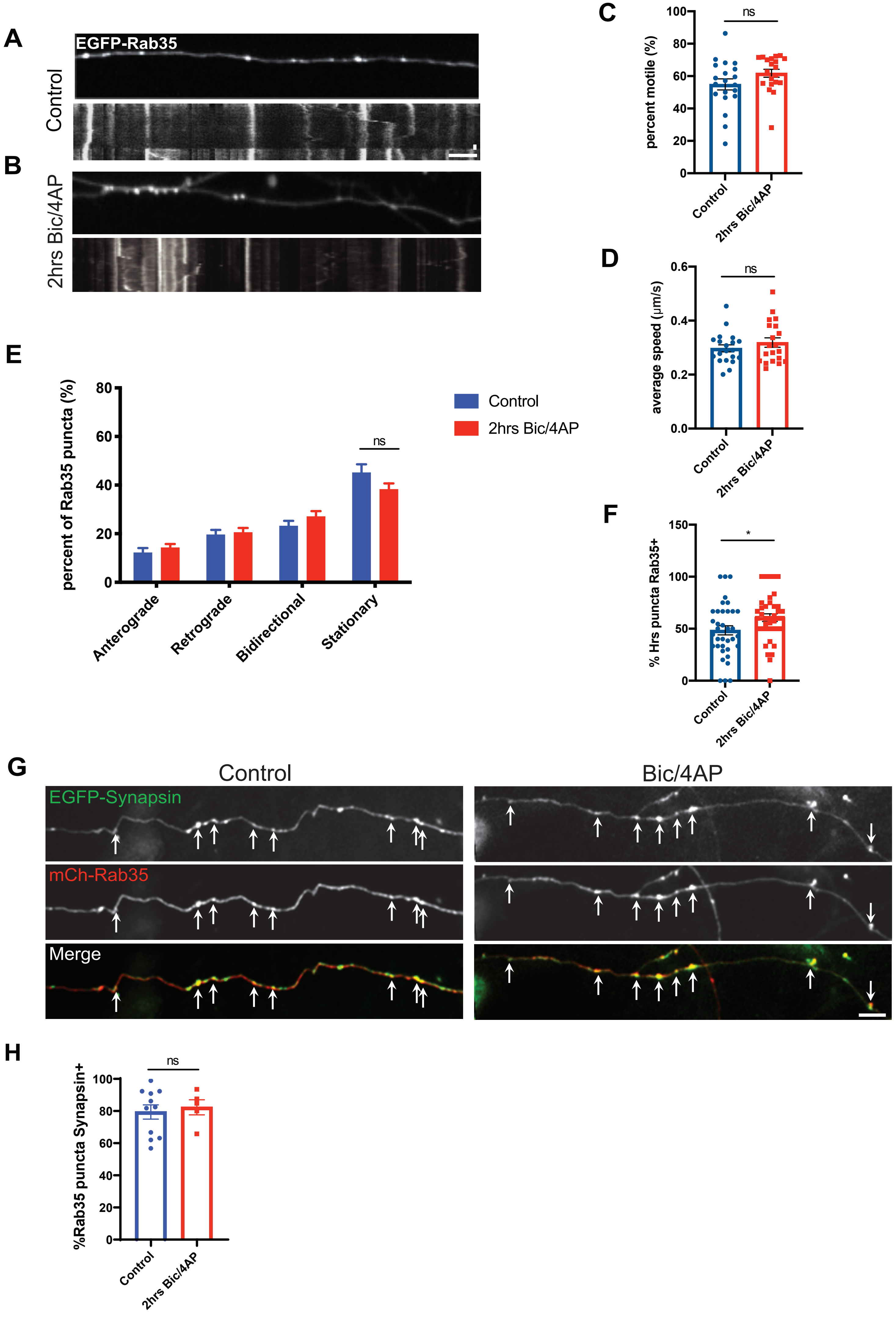

Supplemental Figure 7

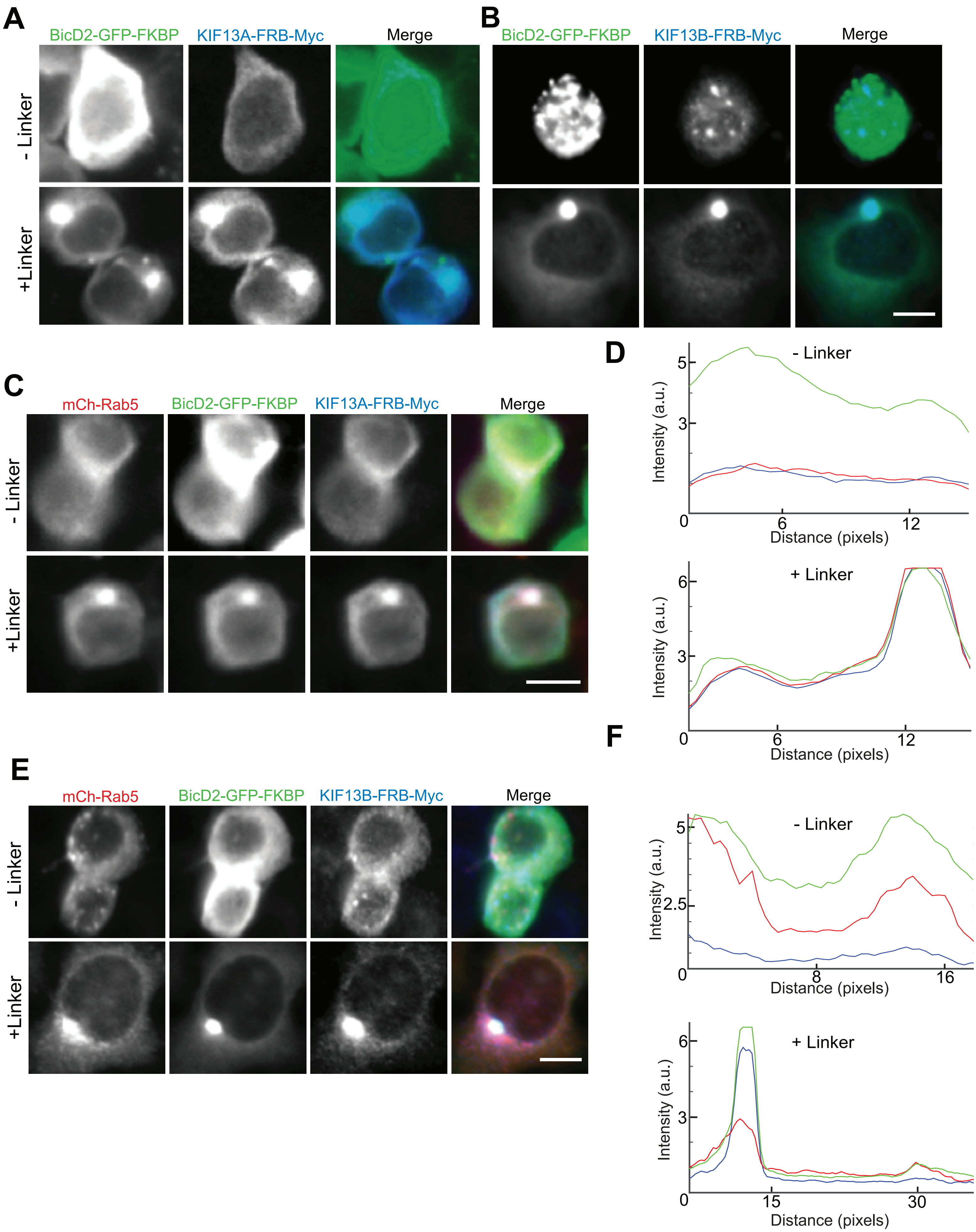

Supplemental Figure 8

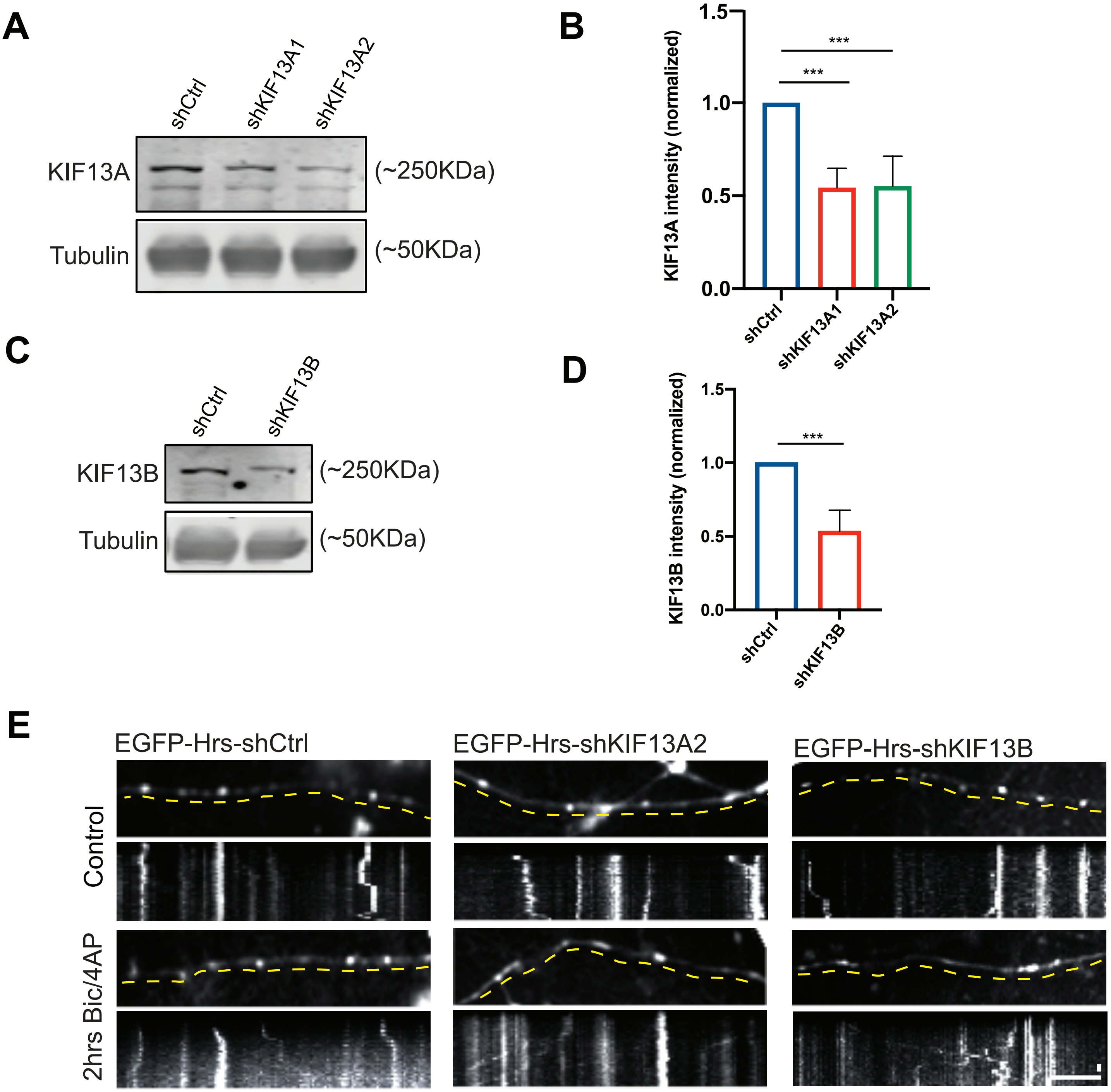
